# Deletion of Nrf1α promotes glutamine addiction of HepG2 cells and thus enhances its apoptosis caused by glutamine deprivation

**DOI:** 10.1101/2025.03.17.643732

**Authors:** Rongzhen Deng, Qun Zhang, Keli Liu, Shaofan Hu, Reziyamu Wufuer, Xi Cheng, Da Lv, Yiguo Zhang

## Abstract

Glutamine metabolism plays a central role in regulating uncontrolled growth of tumor (exhibiting glutamine addiction) by modulating bioenergetics and redox homeostasis, as well as serving as a precursor for biomass synthesis. Thus, limiting its availability is becoming a potential therapeutic strategy. Since Nrf1 is an indispensable determinant of mitochondrial homeostasis through the integration of a multilevel regulatory network for redox balance, no reports on the regulatory role of Nrf1 in glutamine addiction emerged hitherto. In this study, we found that glutamine deprivation leads to mitochondrial morphological and functional damage, reduced GSH levels, increased ROS, and rapid death of Nrf1α-knockout HepG2 cells. A series of further experiments revealed that such rapid death of Nrf1α-deficient cells from glutamine deprivation is caused by different ways. Loss of Nrf1α results in an obvious enhancement of glycolysis and fatty acid synthesis, leading to extensive catabolism of glutamine into α-KG within mitochondria; the α-KG reverses TCA cycles to generate citrate, which exits to the cytosol for fatty acid synthesis. Simultaneously, increased expression of xCT together with decreased expression of GLUL leads to elevated extracellular efflux of glutamate and reduced capacity for glutamine synthesis. These factors intensify glutamine addiction of Nrf1α-null cells, with reducing GSH synthesis and increasing ROS levels due to dysfunctional mitochondria, ultimately leading to enhanced cell death upon glutamine deprivation. Overall, such insights into Nrf1’s role in cancer glutamine metabolism are conducive to developing much specific preventive and therapeutic strategies by precision Nrf1α-targeting anti-tumorigenesis.

## 1. Introduction

Metabolism is the fundamental process underpinning all cellular functions. Unlike normally differentiated cells, cancer cells undergo extensive metabolic reprogramming that encompasses multiple facets, including nutrient acquisition, macromolecule biosynthesis, and energy production. A hallmark of cancer cells is their elevated glucose consumption, characterized by a predominant conversion of glucose into lactic acid even in the presence of oxygen, a phenomenon known as the Warburg effect. Since its discovery, the Warburg effect has established glucose as a pivotal molecule in tumor metabolism research. Researchers have explored the mechanisms by which this effect influences both tumor cells and neighboring cells, particularly through the application of advanced technologies to elucidate intrinsic mechanisms that impact a wide range of cellular activities, from growth and proliferation to apoptosis, invasion, and metastasis. Beyond maintaining high glycolytic rates, cancer cells also exhibit significant alterations in various energy metabolic pathways, including glucose transport, the tricarboxylic acid (TCA) cycle, glutamine metabolism, mitochondrial oxidative phosphorylation, and the pentose phosphate pathway (PPP) [1]. Traditionally, the TCA cycle is viewed as a mechanism for maximizing ATP production from oxidizable substrates, with each turn producing two CO2 molecules. However, due to the Warburg effect, cancer cells exhibit reduced shunting of glucose to mitochondrial metabolism and decreased ATP production. During cell proliferation, the TCA cycle functions as a crucial hub for the interconversion of various metabolic pathways and metabolites. Cancer cells utilize precursors derived from TCA cycle intermediates to synthesize proteins, lipids, and nucleic acids, leading to continuous efflux of intermediates (cataplerosis). To sustain mitochondrial function, tumor cells must compensate for the depletion of TCA cycle intermediates caused by metabolic reprogramming. Glutamine, a non-essential amino acid with amine functional groups, serves as a rich and versatile nutrient that significantly impacts tumor growth. The consumption rate of glutamine in tumor cells far exceeds that of other amino acids. Studies have shown that C-labeled glutamine participates in the TCA cycle in human glioblastoma cells. When the transport of glucose-derived pyruvate to mitochondria is inhibited, glutamine is converted to glutamate via glutaminase (GLS), and GLUD1 activation promotes the conversion of glutamate to α-ketoglutarate (α-KG), thereby facilitating anaerobic supplementation of the TCA cycle. Besides glucose, cancer cells heavily rely on glutamine as a critical energy source and building block for their growth and proliferation. This pronounced dependence on glutamine for TCA cycle support is referred to as glutamine addiction[2].

Glutamine supplementation to the mitochondrial carbon pool provides essential precursors for maintaining mitochondrial membrane potential and synthesizing nucleotides, proteins, and lipids. In normal cells, fatty acid synthesis supplies various substances required for energy storage, membrane biosynthesis, and signal transduction, with most endogenous fatty acids being synthesized from citrate derived from the TCA cycle. Lipid synthesis involves the transfer of mitochondrial citrate into the cytoplasm, where it is converted into oxaloacetate (OAA) and acetyl-CoA, a key precursor for lipid synthesis. Cancer cells within the tumor microenvironment undergo dynamic fluctuations in nutrient availability during tumor progression. The interplay between carcinogenic signal transduction and lipid metabolism, marked by their synthesis and mutual regulation, enables cancer cells to harness lipid metabolic pathways to support rapid proliferation, survival, migration, invasion, and metastasis. However, the Warburg effect limits glucose entry into the TCA cycle, thereby reducing citrate production and hindering its conversion to fatty acids [3]. Under conditions of severe nutrient deficiency and hypoxia, tumor cells engage in de novo fatty acid synthesis to compensate for this deficiency by supplementing the TCA cycle with alpha-ketoglutarate (alpha-KG) derived from glutamine, using citrate from the TCA cycle as an alternative primary carbon source.

Oxidative stress induced by tumor cells during their occurrence, development, metastasis, and anti-tumor processes leads to an increase in reactive oxygen species (ROS)[4, 5]. Low levels of ROS can activate tumorigenic growth signals. However, in rapidly proliferating cancer cells, ROS levels continue to rise. Excessive accumulation of ROS triggers programmed cell death mechanisms in cancer cells, including autophagy, apoptosis, necroptosis, and ferroptosis[6, 7]. Glutamine metabolites in cancer cells directly regulate ROS levels. The primary mechanism by which glutamine metabolism controls ROS levels is through the synthesis of glutathione (γ-glutamyl cysteine + glycine, GSH), a critical intracellular antioxidant that is highly dependent on glutamine. This protects cancer cells from programmed cell death caused by increased oxidative stress, thereby maintaining their survival[8]. In addition to providing nitrogen and carbon for biosynthesis and energy metabolism, glutamine also directly regulates ROS levels. Given that the growth and proliferation of cancer cells are heavily dependent on glutamine, targeting glutamine metabolism has garnered significant attention as a therapeutic strategy[9, 10].

The cap’n’collar-basic leucine zipper (CNC-bZIP) family is one of the most potent eukaryotic transcription factor families that regulate redox homeostasis. Nrf1 is one of the transcription factors with more complex structure and more protein subtypes in this family. It can form heterodimers with sMaf or other bZIP family proteins and bind to antioxidant and/or electrophilic response elements (AREs/EpREs) in the target promoter region to activate downstream gene expression. Studies have shown that Nrf1 is involved in the regulation of antioxidant response, proteasome homeostasis, inflammatory response, lipid metabolism and cell differentiation. Nrf1 is implicated in numerous cellular biological processes, such as the antioxidant response, proteasome homeostasis, inflammatory response, regulation of cell differentiation, DNA damage repair, and cell metabolism. Nrf1 exerts antioxidant and cytoprotective functions by modulating the biosynthesis, utilization, and regeneration of three key antioxidant molecules: glutathione, thioredoxin, and NADPH. As an indispensable regulator of mitochondrial redox homeostasis, the loss of Nrf1 function results in severe mitochondrial dysfunction, leading to potentially fatal oxidative stress. Amplification of reactive oxygen species (ROS) production and glutathione (GSH) depletion in cancer cells represents a promising strategy for disrupting REDOX homeostasis in cancer therapy[11].

High aerobic glycolysis rates in cancer cells are often attributed to mitochondrial dysfunction; however, substantial evidence suggests that the mitochondrial respiratory capacity and fatty acid synthesis in cancer cells are closely linked to glutamine metabolism, particularly under conditions of metabolic stress or carcinogenic activation. Deprivation of glutamine leads to increased mitochondrial reactive oxygen species (ROS) production and decreased GSH levels, ultimately triggering cell death. Metabolic stress is widespread in the tumor microenvironment. Nutrient deficiency directly reduces the production of ATP and leads to excessive production of reactive oxygen species (ROS). ROS accumulation activates multiple pathways and has different effects on cancer cell survival. In this study, we found that glutamine deprivation leads to mitochondrial morphological and functional damage, reduced GSH levels, increased ROS, and rapid cell death in Nrf1α-knockout HepG2 cells. Further investigation revealed that the rapid death of Nrf1α-knockout HepG2 cells under glutamine deprivation is caused by the following factors: After Nrf1α knockout, glycolysis and fatty acid synthesis are enhanced in HepG2 cells, leading to extensive glutamine catabolism into α-ketoglutarate (α-KG) within mitochondria. This α-KG then reverses the TCA cycle to generate citrate, which exits to the cytosol for fatty acid synthesis. Simultaneously, increased expression of xCT and decreased expression of GLUL result in elevated extracellular efflux of glutamate and reduced capacity for glutamine synthesis. These factors intensify glutamine addiction in Nrf1α-knockout HepG2 cells, thereby reducing GSH synthesis and increasing ROS levels due to insufficient mitochondrial function, ultimately leading to cell death under conditions of glutamine deprivation.

## 2. Materials and methods

### 2.1 Cell lines, culture and transfection

The human HepG2 cells (WT) were obtained from the American Type Culture Collection (ATCC, Manassas, VA, USA). Nrf1α-KO cells (cells with Nrf1α gene knockout) were generated through TALEN-mediated genome editing of HepG2 cells[12]. Nrf1α-OE cells (cells with stable Nrf1α overexpression) were primarily constructed using lentiviral technology. All these cell lines were cultured at 37 °C with 5 % CO2 in Dulbecco’s modified Eagle’s medium (DMEM) containing 5mM glutamine (Glutamine deprivation was 0Mm), 25 mmol/L glucose, 10 % (v/v) fetal bovine serum (FBS), 100 units/mL penicillin-streptomycin, before subsequent experiments were performed. In addition, the specified constructs were mixed with Lipofectamine®3000 reagents from Opti-MEM (gibca, Waltham, MA, USA) and some cell lines were transfected for 8 hours. The cells were then allowed to be transfected in a fresh complete culture medium for 24 hours before other experiments were performed.

### 2.2 Plasmid construction

The expression vector for human Nrf1α was constructed by inserting the cDNA sequence of human Nrf1α into the pcDNA3.1/V5His B vector using KpnI and XbaI restriction sites. The expression construct for human GLUL was generated by inserting its cDNA sequence into the same vector backbone, but utilizing KpnI and XhoI restriction sites. For lentiviral expression, the cDNA sequence of human Nrf1α was inserted into the pLJM-EGFP vector at the NheI and EcoR1 sites. The ARE-luc plasmid was created by cloning the promoter region of the GLUL gene into the PGL3-basic vector, resulting in the PGL3-GLUL-ARE construct(Fig.5A).

### 2.3 Nrf1α-overexpressing cell lines were established by lentivirus

The cDNA encoding Nrf1α was cloned into the pLJM1-EGFP vector to generate the expression construct pLJM1-Nrf1α, which was verified by sequencing. Subsequently, 293T cells were transfected with pLJM1-Nrf1α along with the lentiviral packaging plasmids psPAX2 and pMD2G. After a 24-hour recovery period in complete medium containing 10% FBS, the cells were cultured for an additional 24 hours before collecting the supernatant to obtain the lentivirus. HepG2 cells were then infected with the Nrf1α-expressing lentivirus and selected using puromycin resistance. Successfully infected cells (NRF1α-OE) were isolated and stored for further experiments.

### 2.4 Western blotting (WB)

After washing the experimental cells three times with PBS, cell lysis was performed using a denaturing cracking buffer (0.5% SDS, 0.04 mol/L DTT, pH 7.5, supplemented with one tablet of cOmplete protease inhibitor EASYpack per 10 ml of buffer). The harvested proteins were separated by SDS-PAGE on gels containing 8%, 10%, and 15% polyacrylamide. The proteins were then mixed with 3x loading buffer (187.5 mmol/L Tris-HCl, pH 6.8, 6% SDS, 30% glycerol, 150 mmol/L DTT, 0.3% bromophenol blue) to achieve denaturation at 100 °C for 10 minutes. The denatured proteins were subsequently separated by SDS-PAGE on gels containing 8%, 10%, and 15% polyacrylamide, transferred to a polyvinylidene difluoride (PVDF) membrane (Millipore, Billerica, MA, USA), and visualized by western blotting using various antibodies, as shown in the figure. Beta-actin served as an internal control to ensure equal protein loading in each electrophoresis lane.The antibodies used in the study are shown in Table 2.

### 2.5 Real-time qPCR analysis of mRNA expression

Total RNA was isolated using an RNA extraction kit (TIANGEN, Beijing, China), and the extracted RNA was reverse-transcribed into cDNA using a reverse transcription kit (Promega, USA). Real-time quantitative PCR (RT-qPCR) was employed to assess the mRNA expression levels of the target genes in the cells. The primers utilized for RT-qPCR are detailed in Table 1.

### 2.6 Dual-luciferase reporter gene assays

An equal number of 1.0 × 105 COS-1 cells were seeded in 12-well plates and allowed to grow until they reached 70-80% confluence. Transfection was then performed using Lipofectamine®3000 reagent in Opti-MEM (Gibco, Waltham, MA, USA) for a total duration of 8 hours. The plasmids used for transfection included the pcDNA3.1-Nrf1α vector (or empty pcDNA3.1 vector) expressing Nrf1α, PGL3-GLUL-ARE, and pRL-TK plasmid at a ratio of 5:4:1 (totaling 1.0 μg of transfected plasmid). Following transfection, cells were lysed and cultured in fresh complete medium for an additional 24 hours. Luciferase activity was subsequently measured using the dual luciferase reporter assay system (Promega, Madison, WI, USA).

### 2.7 Intracellular ROS and GSH detection

The experimental cells were incubated with 20,70-Dichlorodihydrofluorescein diacetate (DCFH-DA, S0033, Beyotime, Shanghai, China),Dihydroethidium(DHE, HY-D0079, MedChemExpress),and Monochlorobimane(mBCL, HY-101899, MedChemExpress) in serum-free medium at 37 °C for 20 minutes. Following three washes with fresh serum-free medium, the fluorescence intensity was measured using flow cytometry or inverted fluorescence microscopy under specific excitation (Ex) and emission (Em) wavelengths. Specifically, the Ex/Em wavelengths for DCFH-DA are 488/525 nm, for DHE are 518/610 nm, and for monochlorobimane are 380/470 nm.

### 2.8 pyruvate, Lactic, triglyceride, NADP/NADPH and ATP content detection

After preparing the cells, the culture medium was aspirated and the cells were washed three times with PBS. Subsequently, the levels of ATP (Beyotime, Shanghai, China), pyruvate (Solarbio, Beijing, China), and glucose (Applygen, Beijing, China) in the specified cell line were measured using their respective assay kits. Additionally, intracellular levels of lactate (Solarbio, Beijing, China), triglycerides (Applygen, Beijing, China), and NADP/NADPH (Beyotime, Shanghai, China) were determined. all results were normalized to protein concentration for further analysis.

### 2.9 Lipid droplets stained with oil red O in cells

When the cells reached the desired state, they were fixed using 4% paraformaldehyde (Boster Biological Technology, Wuhan, China) for 15 minutes at room temperature. Subsequently, the cells were stained with a 3g/L Oil Red O solution (Sangon Biotech, Shanghai, China) for 30 minutes. After staining, the cells were washed three times with 60% isopropyl alcohol (Kelong, Chengdu, China) to remove excess stain. Finally, the stained cells were examined under a microscope and images were captured.

### 2.10 Observation of mitochondria by Transmission electron microscopy

When the cells were ready, they were gently scraped off using a cell scraper and subsequently fixed with 2.5% glutaraldehyde in 0.1 M sodium cacodylate buffer (pH 7.4) at 37 °C for 30 minutes. The fixed cells were stored in 0.1 M sodium cacodylate buffer at 4 °C and then incubated at 4 °C for 1 hour with a mixture of 1% osmium tetroxide (OsO4) and 1.5% potassium ferricyanide [K4Fe(CN)6] in 0.1 M sodium cacodylate buffer (pH 7.4). Following this, the samples were incubated overnight at 4 °C in 0.25% uranyl acetate. After three washes, the samples underwent dehydration in a graded ethanol series (25%, 50%, 75%, 95%, and 100% v/v) for 15 minutes each. Subsequently, the samples were infiltrated with 812 Embed resin for 5-8 hours. Ultrathin sections (60-80 nm) were prepared using an ultramicrotome. The sections were stained with a saturated solution of 2% uranyl acetate in ethanol for 8 minutes in the dark, followed by three washes with 70% ethanol. They were then washed three times with ultra-pure water and stained with 2.6% lead citrate for 8 minutes in a CO2-free environment. Finally, the sections were rinsed three more times with ultra-pure water and dried on filter paper. Transmission electron microscopy was used to analyze the images.

### 2.11 Flow cytometry analysis of apoptosis and JC-1

The cells were trypsinized at 1000g for 5 minutes and subsequently centrifuged. Following this, the cells were washed three times with PBS. JC-1 and V-FITC were diluted in serum-free DMEM medium and incubated with the cells for 20 minutes.The results were analyzed by the FlowJo 7.6.1 software (FlowJo, Ashland, OR, USA) before being presented.

### 2.12 Immunocytochemistry and confocal microscopy

After being disinfected by soaking in alcohol, the glass cover slide (1 cm²) was placed flat in a 6-well plate. Cells were inoculated onto the glass cover slide and cultured for 24 hours at room temperature. Subsequently, the cells were fixed with polyformaldehyde fixative (Servicebio, Wuhan, China) for 30 minutes at room temperature, followed by permeabilization with 0.2% Triton X-100 for 20 minutes. The fixed cells were incubated with 1% BSA for 60 minutes to block non-specific binding sites. Primary antibodies against Nrf1 and GLUL were then added and incubated overnight at 4°C. After washing five times with PBS, the cells were incubated with Alexa Fluor 488-conjugated Goat anti-mouse IgG (H + L) or TRITC-conjugated Goat Anti-Rabbit IgG (H + L) (both from ZSBG-Bio, Beijing, China) in the dark at room temperature for 4 hours. The cells were gently washed with PBS five times, followed by nuclear staining with DAPI (Beyotime, C1005). Immunofluorescence images were captured using confocal microscopy.

### 2.13 Metabolomics analysis

Cells were rapidly frozen in liquid nitrogen after culturing and subjected to untargeted metabolomics analysis (with at least three replicates per group), with technical support provided by Wuhan Maiwei Biotechnology Co., Ltd. After obtaining the raw data, the fold change in expression levels (Log2 FC) was calculated using software tools such as R, Excel, and Notepad++. The Human Metabolome Database (HMDB) and KEGG database were then used to enrich or mine for differentially expressed metabolites in cell and tissue samples. Differences in metabolites and enriched pathways were visualized using R, MetaboAnalyst, Canvas X, and other online tools and software.

### 2.14 Statistical analysis

Significant differences in the obtained data were statistically assessed using Student’s t-test or multivariate analysis of variance (MANOVA). The results are expressed as fold changes (mean ± standard deviation), with each datum representing at least three independent experiments, each conducted in triplicate.

## 3. Results

### 3.1 Glutamine deprivation results in the rapid demise of Nrf1α gene knockout cells

Glutamine, a non-essential amino acid abundant in human plasma, becomes conditionally essential in rapidly dividing cancer cells due to their increased demand. This phenomenon, known as glutamine addiction, has been extensively studied, revealing that many tumor cell types fail to proliferate in the absence of glutamine. In mammalian cells, Nrf1, a key member of the CNC-bZIP family, regulates oxidative stress response genes through ARE-binding sequences in promoter regions, thereby influencing tumor progression. To investigate the expression of Nrf1α in hepatocellular carcinoma (HCC) cells under glutamine deprivation and assess their sensitivity to such conditions, we examined the protein and mRNA levels of Nrf1α in HepG2, Huh7, and MHCC97H cells at various time points following glutamine starvation using Western blot and RT-PCR. The results indicated that Nrf1α protein expression initially increased and then decreased in all three cell lines after glutamine deprivation. Specifically, Nrf1α protein levels peaked at 24 hours post-glutamine starvation in HepG2 and MHCC97H cells before declining. In contrast, Nrf1α protein expression in Huh7 cells reached its peak at 36 hours and subsequently decreased.

To investigate the relationship between glutamine deprivation and Nrf1α in hepatocellular carcinoma cells, a stable Nrf1α -overexpressing cell line (NRF1α-OE) was generated by infecting HepG2 cells with lentivirus carrying the Nrf1α gene. Concurrently, glutamine deprivation experiments were performed using an Nrf1α-knockout HepG2 cell line (Nrf1α-KO) that was previously established in our laboratory. The successful generation of these cell lines was confirmed through western blotting and real-time qPCR analysis(Fig. 1, D). Compared with wild-type (WT) cells, western blot and real-time qPCR analyses revealed that Nrf1α-KO cells exhibited negligible expression of the Nrf1α protein (Fig. 1D,d1), along with a marked reduction in mRNA levels. Conversely (Fig. 1D,d3), Nrf1α-OE cells demonstrated a significant increase in both Nrf1α protein(Fig. 1D,d1) abundance and basal mRNA expression(Fig. 1D,d3).

**Fig. 1.**
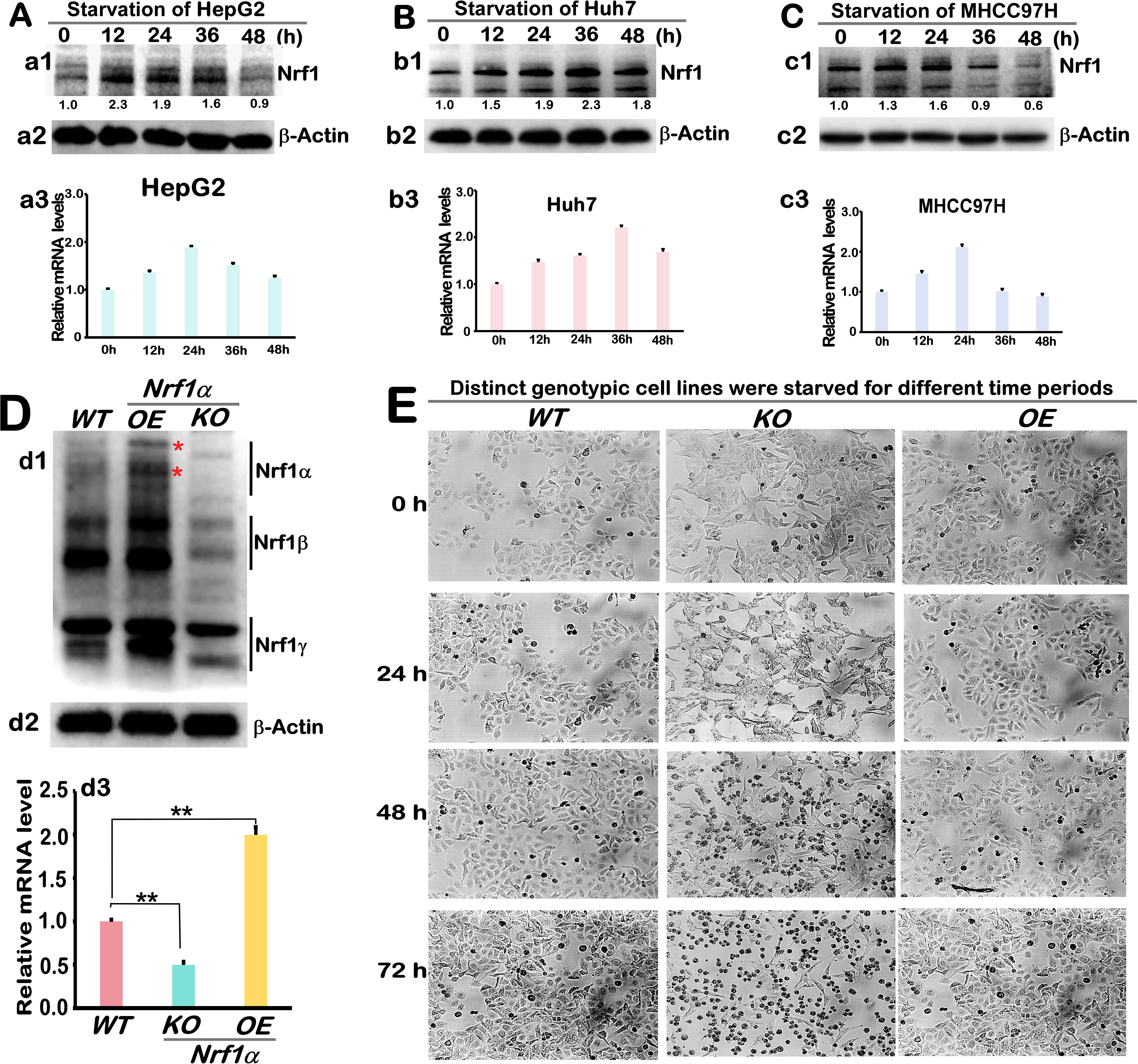
Glutamine deprivation led to extensive mortality in Nrf1α-KO cells. A. Nrf1α protein (a1) and Nrf1 mRNA (a3) levels were assessed by Western blotting and RT-qPCR at 0h, 12h, 24h, 36h, and 48h following glutamine deprivation in HepG2 cells. Beta-actin served as the loading control, The results are presented as mean ± SEM (n=3x3). B. Nrf1α protein (a1) and Nrf1 mRNA (a3) levels were assessed by Western blotting and RT-qPCR at 0h, 12h, 24h, 36h, and 48h following glutamine deprivation in Huh7 cells. Beta-actin served as the loading control, The results are presented as mean ± SEM (n=3x3). C. Nrf1α protein (a1) and Nrf1 mRNA (a3) levels were assessed by Western blotting and RT-qPCR at 0h, 12h, 24h, 36h, and 48h following glutamine deprivation in MHCC97H cells. Beta-actin served as the loading control, The results are presented as mean ± SEM (n=3x3). D. Nrf1 protein levels in three genotypic cell lines (WT, Nrf1α-KO, and Nrf1α-OE) were assessed by western blotting with specific antibodies. Beta-actin served as the loading control(d1). Nrf1 mRNA levels in three genotypic cell lines (WT, Nrf1α-KO, and Nrf1α-OE) were assessed by RT-qPCR (d2). The results are presented as mean ± SEM (n=3x3; *, p < 0.05 and **, p < 0.01). E. The morphological changes of glutamine-deprived WT, Nrf1α-KO, and Nrf1α-OE cell lines were observed under a microscope at 0h, 24h, 48h, and 72h (with an original magnification of 200x).

Cell morphology was monitored at various time points during glutamine starvation. The results demonstrated that all three cell lines exhibited normal growth and morphology at the 0-hour time point. After 12 hours of glutamine deprivation, the Nrf1α-KO cell line showed altered morphology, with cells becoming elongated and their edges and contours becoming less defined. In contrast, both the wild-type (WT) strain and the Nrf1α-overexpressing (Nrf1α-OE) cell line maintained normal growth without significant morphological changes. By 48 hours of glutamine starvation, the Nrf1α-KO cell line displayed clear signs of cell death, whereas the WT and NRF1-OE cell lines remained unaffected. At 72 hours, a substantial proportion of the Nrf1α-KO cells had died, while the WT and Nrf1α-OE cell lines continued to exhibit no significant abnormalities. These findings suggest that Nrf1α-KO cells are highly sensitive to glutamine deprivation(Fig. 1E).

### 3.2 Glutamine deprivation leads to Nrf1α-deficient cell mitochondrial dysfunction and apoptosis

Based on the aforementioned experiments, we conducted DCFH staining flow cytometry analysis on three cell types: WT, Nrf1α-KO, and Nrf1α-OE, which were cultured under glutamine starvation for 24 hours. The results indicated that glutamine deprivation induced substantial apoptosis and necrosis in Nrf1α-KO cells, further confirming confirmed that glutamine deprivation can cause rapid death of Nrf1α-KO cells (Fig. 2A).

**Fig. 2.**
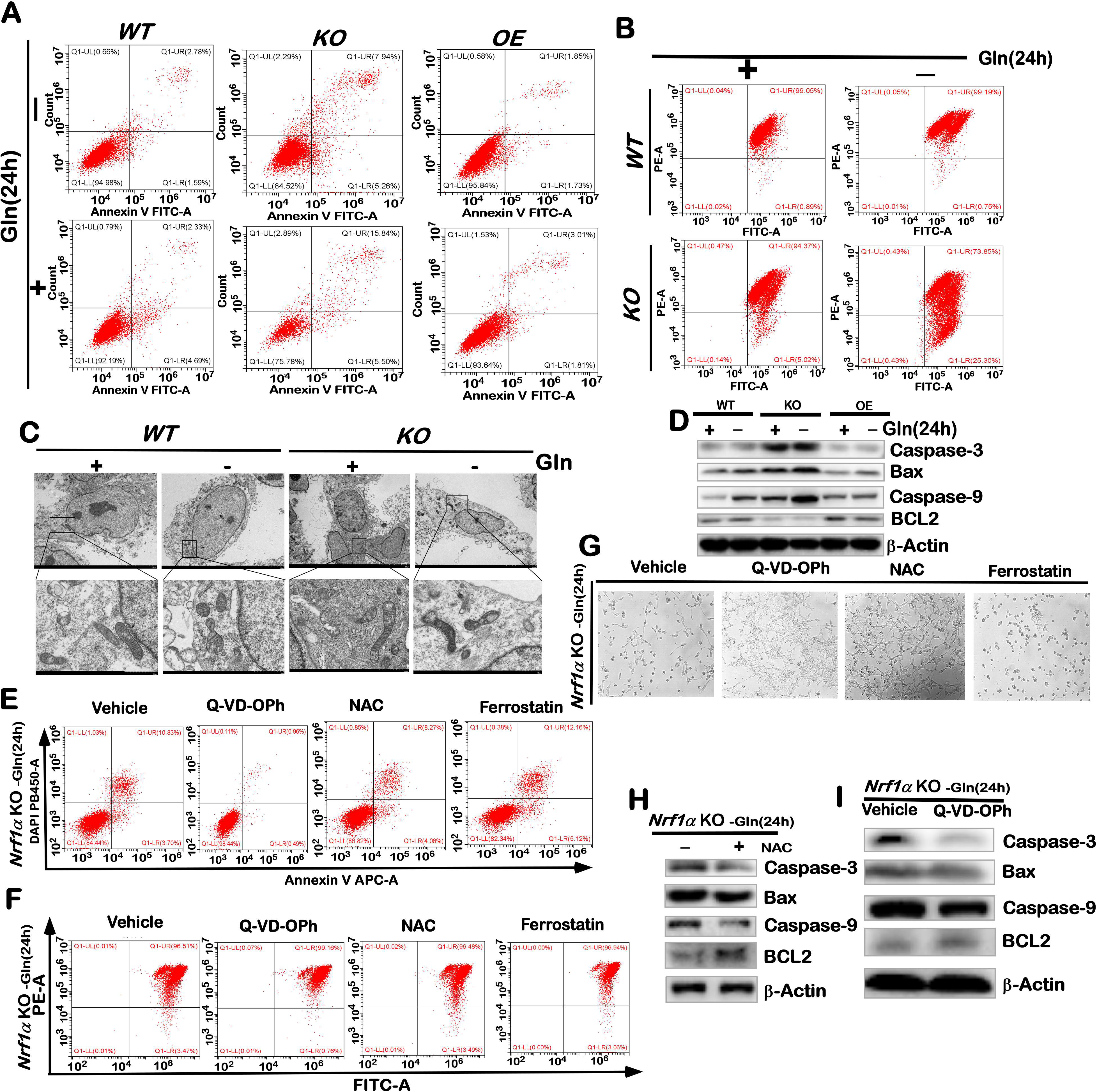
Glutamine deprivation results in mitochondrial morphological and functional abnormalities and accelerated cell apoptosis in Nrf1α-KO cells. A. The apoptosis of glutamine-deprived cells in WT, Nrf1α-KO, and Nrf1α-OE cell lines was analyzed by flow cytometry following incubation with Annexin V-FITC and PI. B. Mitochondrial membrane potential in glutamine-deprived WT and Nrf1α-KO cells was assessed using JC-1 staining and flow cytometry. C. Transmission electron microscopy was employed to examine the morphology of glutamine-deprived WT and Nrf1α-KO cells and their mitochondria. D. Western blotting was used to detect the expression levels of Caspase-3, Caspase-9, Bax, and BCL2 proteins in WT, Nrf1α-KO, and Nrf1α-OE cell lines after glutamine deprivation. E. Nrf1α-KO cells were cultured for 24 hours in glutamine-free medium supplemented with q-VD-OPH (10 μM), NAC (10 μM), and Ferrostatin-1 (2 μM). Apoptosis was then analyzed by flow cytometry after staining with Annexin V-FITC and PI. F. Nrf1α-KO cells were cultured for 24 hours in glutamine-free medium containing q-VD-OPH (10 μM), NAC (10 μM), and Ferrostatin-1 (2 μM), followed by mitochondrial membrane potential analysis using JC-1 staining and flow cytometry. G. The morphological changes in Nrf1α-KO cells were observed under a microscope after 48 hours of culture in glutamine-free medium supplemented with q-VD-OPH (10 μM), NAC (10 μM), and Ferrostatin-1 (2 μM), with an original magnification of 200x H. Western blotting was performed to detect the expression of Caspase-3, Caspase-9, Bax, and BCL2 proteins in Nrf1α-KO cells cultured for 24 hours in glutamine-free medium containing NAC (10 μM). I. Western blotting was also used to analyze the expression of Caspase-3, Caspase-9, Bax, and BCL2 proteins in Nrf1α-KO cells cultured for 24 hours in glutamine-free medium supplemented with q-VD-OPH (10 μM).

Mitochondria play a critical role in various biochemical processes, and cell death is often linked to the disruption of mitochondrial transmembrane potential. The mitochondrial membrane potential was typically assessed using the JC-1 probe. JC-1 results indicated that the membrane potential of WT cells remained largely stable after 24 hours of glutamine deprivation. In contrast, the mitochondrial membrane potential of Nrf1α-KO cells showed a slight decrease compared to wild-type cells under normal conditions, but it significantly declined following 24 hours of glutamine starvation. These findings suggest that glutamine deprivation induces mitochondrial dysfunction in Nrf1α gene knockout cell lines (Fig. 2B). Mitochondria are highly dynamic intracellular organelles characterized by ultrastructural heterogeneity, reflecting cellular behavior and function. During metabolic or environmental stress, mitochondrial fission and fusion play a pivotal role in maintaining mitochondrial function, enabling these organelles to respond to diverse cellular needs and signals. This process is also a critical component of the mechanism by which mitochondria adapt to pathophysiological conditions that challenge cellular homeostasis. Mitochondrial fusion integrates damaged mitochondria, followed by their separation through mitochondrial division and subsequent removal via autophagy. However, if the stress persists, it may overwhelm the cell’s adaptive capacity, leading to defective changes in mitochondrial morphology (indicating impaired mitochondrial function or the onset of mitochondrial disease). We examined the mitochondrial morphology of WT and Nrf1α-KO cells under normal medium conditions and after 24 hours of glutamine deprivation using transmission electron microscopy. The results revealed that most mitochondria in the WT group cultured in normal medium were elliptical and rod-shaped, with clearly visible cristae (Fig. 2C).

Nrf1 and glutamine play crucial roles in regulating cellular energy metabolism and redox homeostasis. To determine whether the rapid death of Nrf1α-KO cells following glutamine deprivation is due to apoptosis or necrosis, we selected several cell death inhibitors to investigate their effects on cell viability. These inhibitors included the caspase inhibitor Q-VD-OPH, the ferroptosis inhibitor Ferrostatin-1, and the potent ROS scavenger N-acetyl-L-cysteine (NAC). After 24 hours of treatment with these inhibitors, apoptosis was assessed using DCFH staining, and mitochondrial membrane potential was evaluated using JC-1 staining. The results indicated that Q-VD-OPH and NAC could mitigate apoptosis induced by glutamine deprivation, whereas Ferrostatin-1 unexpectedly promoted apoptosis rather than alleviating it (Fig. 2E). JC-1 staining revealed that Q-VD-OPH rescued the decreased mitochondrial membrane potential in Nrf1α-KO cells caused by glutamine deprivation. However, NAC and Ferrostatin-1 did not significantly protect against the reduced membrane potential induced by glutamine deprivation (Fig. 2F). When the duration of glutamine starvation was extended to 48 hours, Q-VD-OPH and antioxidants demonstrated a significant cytoprotective effect on cell death caused by glutamine deficiency, leading to increased cell survival. In contrast, Ferrostatin-1 not only failed to protect against cell death but also promoted it (Fig. 2G). These findings suggest that the extensive cell death induced by Nrf1α-KO and glutamine deprivation may be attributed to mitochondrial dysfunction and apoptosis resulting from intracellular redox imbalance.

To further confirm whether the massive death of Nrf1α-KO cells induced by glutamine deprivation is due to apoptosis, we experimentally detected the expression levels of apoptosis-related genes including Caspase-3, Caspase-9, Bax, and BCL2. Western blot(WB) and real-time quantitative PCR (RT-qPCR) analysis revealed that compared with the WT group, the expression levels of Caspase-3, Caspase-9, and Bax were significantly increased in Nrf1α-KO cells, while BCL2 expression was decreased. In contrast, in Nrf1α-OE cells, Caspase-3 expression remained unchanged, whereas Caspase-9 and BCL2 expression increased, and Bax expression decreased(Fig. 2D and Fig.S2 A). After 24 hours of glutamine deprivation, Caspase-9 and Bax expression levels were significantly elevated in all three cell lines, while Caspase-9 and BCL2 expression did not change(Fig. 2D and Fig.S2 A). Additionally, we examined the expression of apoptosis-related genes in Nrf1α-KO cells after adding Q-VD-OPH and NAC to the glutamine medium. The results showed that the expression of Caspase-9, Caspase-3, and Bax decreased following Q-VD-OPH treatment, while BCL2 expression remained unchanged(Fig. 2I). In the NAC-treated group, the protein levels of Caspase-9, Caspase-3, and Bax were significantly reduced, while BCL2 expression was significantly increased(Fig. 2H). These findings suggest that the rapid death of Nrf1α-KO cells under glutamine deficiency is likely caused by an imbalance in cellular redox status.

### 3.3 Glutamine deprivation results in inadequate GSH synthesis in Nrf1α-deficient cell, leading to cellular redox imbalance and ultimately apoptosis

Redox reactions involve the transfer of electrons from a reducing agent to an acceptor molecule, ultimately leading to the generation of reactive oxygen species (ROS).The experiment detected the ROS levels of WT and Nrf1α-KO after glutamine deprivation through two ROS probes, namely DCFH-DA (which can detect substances ranging from hydrogen peroxide (H₂O₂), superoxide anion (O₂⁻), hydroxyl radical (·OH) to peroxynitrite (ONOO⁻)) and Dihydroethidium (DHE) (mainly used for detecting superoxide anion).Compared with WT cells, Nrf1α-KO cells exhibited significantly higher ROS levels. Both WT and Nrf1α-KO cells showed a significant increase in ROS levels after 24 hours of glutamine deprivation(Fig. 3A). To validate these findings, the experiment was repeated using a DHE probe, and fluorescence intensity was observed under an inverted fluorescence microscope. The results were consistent with those obtained using the DCFH-DA probe(Fig. 3B). Additionally, ROS levels were measured in glutamine-deprived Nrf1α-KO cells treated with Q-VD-OPH, Ferrostatin 1, and NAC. The results indicated that Ferrostatin 1 increased cellular ROS levels, whereas NAC reduced them(Fig. 3H). These findings suggest that Nrf1α deficiency in HepG2 cells leads to severe endogenous oxidative stress, which is exacerbated by glutamine deprivation.

**Fig. 3.**
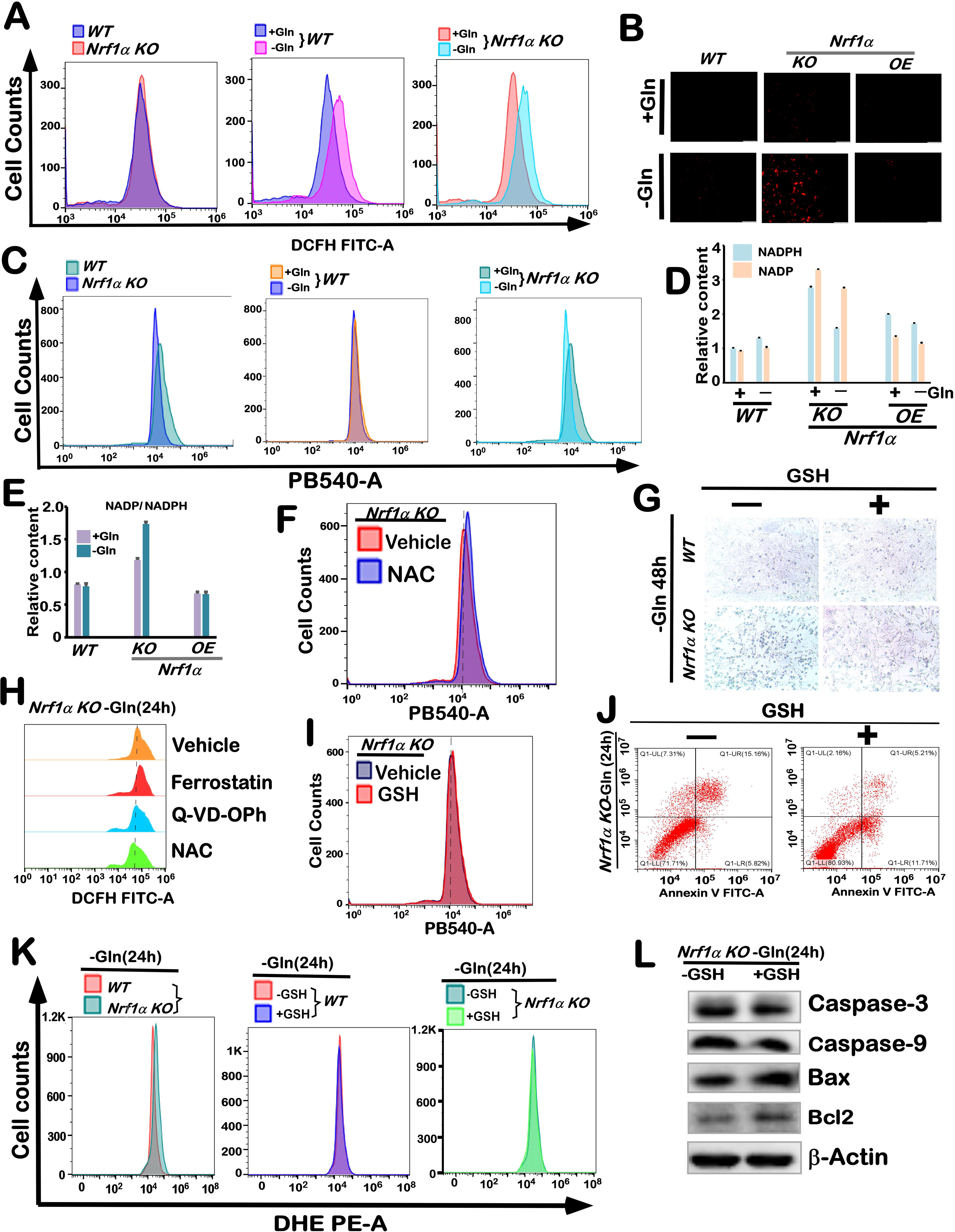
Glutamine deprivation leads to a reduction in the GSH content of Nrf1α gene knockout cells, consequently inducing excessive oxidative stress and apoptosis. A. WT and Nrf1α-KO cells were subjected to glutamine deprivation for 24 hours, followed by staining with DCFH for 30 minutes. Changes in reactive oxygen species (ROS) levels were subsequently analyzed using flow cytometry. B. After 24 hours of glutamine deprivation, WT, Nrf1α-KO, and Nrf1α-OE cells were stained with DHE for 30 minutes, and ROS changes were observed using fluorescence microscopy. C. Following 24 hours of glutamine deprivation and 30 minutes of mBCL staining, the changes in GSH levels in WT and Nrf1α-KO cells were detected by flow cytometry. D. The NADPH and NADP levels in WT, Nrf1α-KO, and Nrf1α-OE cells after 24 hours of glutamine deprivation were measured using a detection kit. E. The ratio of NADP/NADPH in WT, Nrf1α-KO, and Nrf1α-OE cells after 24 hours of glutamine deprivation was determined using a detection kit. F. Nrf1α-KO cells were cultured for 24 hours in glutamine-free medium containing NAC (10 μM) and stained with mBCL for 30 minutes. Changes in GSH levels were detected by flow cytometry. G. The morphology of Nrf1α-KO cells was observed under a microscope after 48 hours of culture in glutamine-free medium supplemented with GSH (5 mM). (with an original magnification of 200x) H. WT, Nrf1α-KO, and Nrf1α-OE cells were cultured for 24 hours in glutamine-free medium containing q-VD-OPH (10 μM), NAC (10 μM), and Ferrostatin-1 (2μM), respectively, and stained with DCFH for 30 minutes. Changes in ROS levels were detected by flow cytometry. I. Nrf1α-KO cells were cultured for 24 hours in glutamine-free medium containing GSH (5 mM) and stained with mBCL for 30 minutes. Changes in GSH levels were detected by flow cytometry. J. Nrf1α-KO cells were cultured for 24 hours in glutamine-free medium containing GSH (5 mM) and analyzed for apoptosis using Annexin V-FITC and PI staining by flow cytometry. K. WT and Nrf1α-KO cells were cultured for 24 hours in glutamine-free medium containing GSH (5 mM) and stained with mBCL for 30 minutes. Changes in GSH levels were detected by flow cytometry. L. The expression of Caspase-3, Caspase-9, Bax, and Bcl-2 proteins in Nrf1α-KO cells cultured for 24 hours in glutamine-free medium containing GSH (5 mM) was detected by Western blotting.

Reduced nicotinamide adenine dinucleotide phosphate (NAD(P)H and glutathione (GSH) are essential for maintaining cellular REDOX homeostasis and regulating cellular metabolism[13, 14]. To investigate the cause of increased ROS levels following glutamine deprivation in NRF1-α-deficient cells, we measured the NADP, NADPH, and NADP/NADPH ratios using a commercially available kit. The results demonstrated that both NRF1-α knockout (KO) and overexpression (OE) significantly increased the levels of NADP and NADPH(Fig. 3D). However, the NADP/NADPH ratio was significantly elevated in NRF1-α KO cells but slightly decreased in NRF1-α OE cells(Fig. 3E). Upon glutamine [deprivation, the NADP and NADPH levels and the NADP/NADPH ratio were significantly increased in NRF1-α KO cells, whereas WT and NRF1-α OE cells exhibited decreased NADP and NADPH levels with little change in the NADP/NADPH ratio(Fig. 3D-3E). These findings further confirm that glutamine deprivation exacerbates oxidative stress in NRF1-α-deficient cells.

Simultaneously, the intracellular GSH levels were measured using the mBCL probe. The experimental results demonstrated that, compared to the control group, GSH levels were elevated in Nrf1α-KO cells. However, upon glutamine deprivation, GSH levels in Nrf1α-KO cells markedly decreased. In contrast, GSH levels in WT cells remained relatively stable following glutamine deprivation(Fig. 3C). Glutamine metabolism regulates oxidative homeostasis through multiple pathways, including the biosynthesis of glutathione (GSH)[15]. GSH serves as a reducing agent with various physiological functions, such as scavenging free radicals, providing antioxidant protection, and neutralizing electrophiles. Disruption of the ROS/GSH balance leads to harmful oxidation and chemical modification of biomacromolecules, ultimately resulting in cell cycle arrest, inhibition of proliferation, and potentially induced cell death[16]. Therefore, we hypothesized that the apoptosis of glutamine-deprived Nrf1α-KO cells may be attributed to the significant reduction in GSH levels. To test this hypothesis, we experimentally treated Nrf1α-KO cells with glutamine deprivation while supplementing GSH and NAC. The results demonstrated that after 24 hours of treatment with GSH and NAC, the intracellular GSH content increased(Fig.3F and 3I), and ROS (Fig.3K) levels in Nrf1α-KO cells were alleviated following glutamine deprivation supplemented with GSH. Moreover, GSH supplementation significantly mitigated apoptosis in Nrf1α-KO cells induced by glutamine deprivation(Fig.3J). We extended the GSH treatment duration to 48 hours and observed cell morphology under a microscope. After 48 hours, it was evident that the control group of Nrf1α-KO cells subjected to glutamine deprivation exhibited substantial cell death, whereas the addition of GSH effectively reduced cell death caused by glutamine deprivation(Fig.3G). To further elucidate the underlying mechanisms, we examined the protein expression of key apoptotic genes via Western blotting. The findings indicated that GSH supplementation significantly decreased the expression of pro-apoptotic proteins Caspase-9 and Caspase-3 in Nrf1α-KO cells induced by glutamine deprivation. Additionally, BCL2 protein expression was markedly increased, while Bax gene expression remained unchanged(Fig.3L). Based on these experimental results, it can be inferred that the extensive cell death following glutamine deprivation is primarily due to apoptosis triggered by REDOX imbalance resulting from decreased GSH levels.

### 3.4 Nrf1α gene knockout resulted in enhanced glycolysis and lipid anabolism, leading to increased lipid deposition. However, glutamine deprivation mitigated cellular lipid accumulation while concurrently disrupting mitochondrial function

As previously mentioned, glutamine deprivation led to a decrease in mitochondrial membrane potential and alterations in mitochondrial morphology in Nrf1α-KO cells. Mitochondria serve as the hub for various metabolic pathways, with their TCA cycle linking glucose catabolism to mitochondrial oxidation(Fig.S1). Nutrients are metabolized and transported into the tricarboxylic acid (TCA) cycle, producing metabolic precursors for large molecules such as ATP, lipids, proteins, DNA, and RNA (Fig.S1). Mitochondria function as bioenergetic power stations, biosynthesis centers, redox balancers, and waste management hubs[17, 18]. Therefore, we investigated the expression of genes related to glucose metabolism, lipid metabolism, the TCA cycle, and changes in metabolites. RT-qPCR and WB results demonstrated that during glycolysis, the expression levels of hexokinase (HK1/HK2) and glucose transporter (SLC2A1/4) genes were significantly elevated in Nrf1α-KO cells but decreased in Nrf1α-OE cells. The expression of HK1, HK2, and SLC2A1/4 genes was reduced following glutamine deprivation(Fig.4A and Fig. S2 B). The phosphofructokinase (PFKL) gene expression was decreased in both Nrf1α-OE and Nrf1α-KO cells, and this reduction was also observed after glutamine starvation. The expression of pyruvate kinase (PKM) and lactate dehydrogenase (LDHA/D) genes was decreased in Nrf1α-KO cells, while no significant changes were noted in Nrf1α-OE cells.

**Fig. 4.**
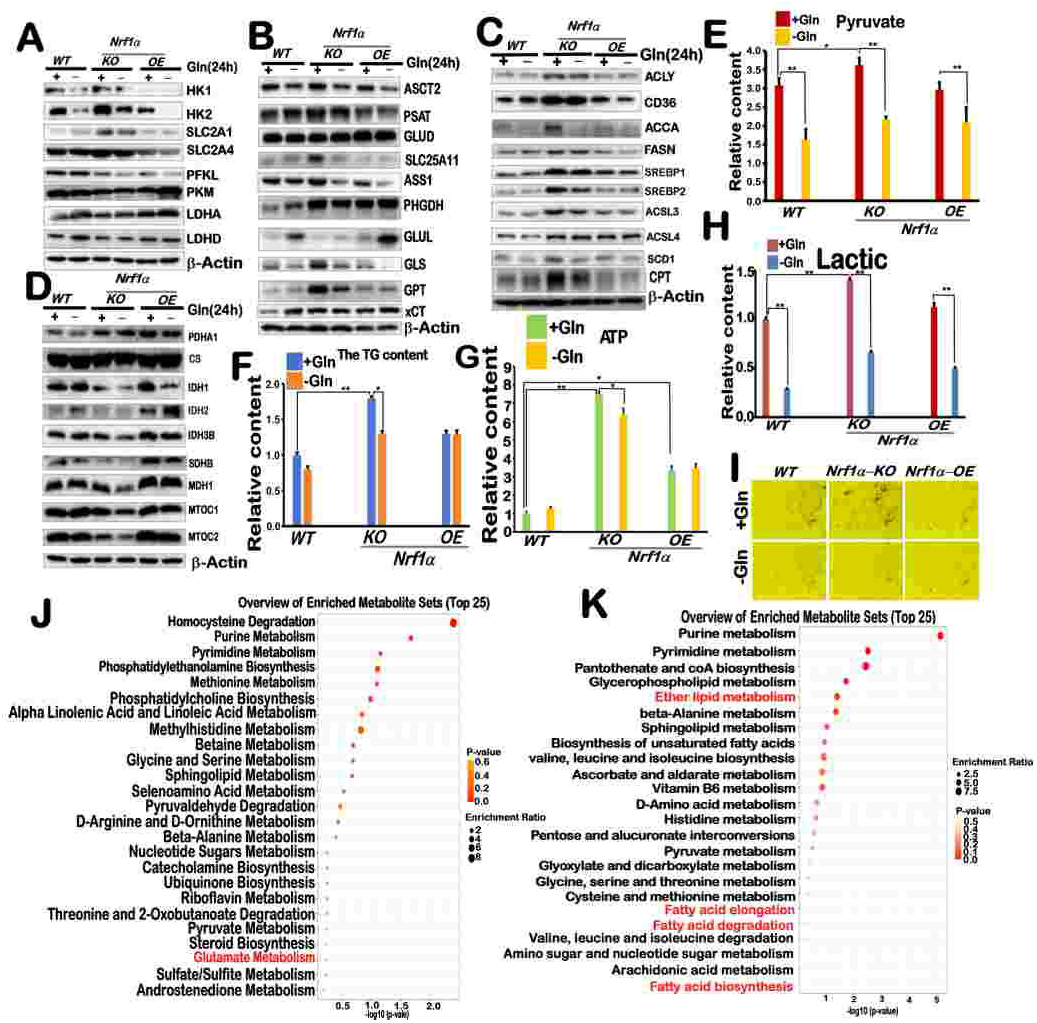
Nrf1α gene knockout resulted in enhanced glycolysis and lipid anabolism, leading to increased lipid deposition. However, glutamine deprivation mitigated cellular lipid accumulation while concurrently disrupting mitochondrial function. A. Western blotting was employed to assess the expression levels of key glucose metabolism proteins, including HK1, HK2, SLC2A1, SLC2A4, PFKL, PKM, LDHA, and LDHD, in WT, Nrf1α-KO, and Nrf1α-OE cells following 24 hours of glutamine deprivation. B. Detection of ASCT2, PSAT, GLUD, SLC25A11, ASS1, PHGDH, GLUL, GLS, GPT, and xCT protein expression related to glutamine metabolism in WT, Nrf1α-KO, and Nrf1α-OE cells after 24 hours of glutamine deprivation by Western blotting. C. Detection of ACLY, CD36, ACCA, FASN, SREBP1, SREBP2, ACSL3, ACSL4, SCD1, and CPT protein expression related to lipid metabolism in WT, Nrf1α-KO, and Nrf1α-OE cells after 24 hours of glutamine deprivation by Western blotting. D. Detection of PDHA1, CS, IDH1, IDH2, IDH3B, SDHB, MDH1, MTOC1, and MTOC2 protein expression related to the TCA cycle in WT, Nrf1α-KO, and Nrf1α-OE cells after 24 hours of glutamine deprivation by Western blotting. E. Measurement of pyruvate levels in WT, Nrf1α-KO, and Nrf1α-OE cells following glutamine deprivation using a kit. The data are shown as fold changes (the mean ± SEM, n = 3 × 3; *, p < 0.05 and **, p < 0.01). F. Measurement of triglyceride (TG) content in WT, Nrf1α-KO, and Nrf1α-OE cells following glutamine deprivation using a kit. The data are shown as fold changes (the mean ± SEM, n = 3 × 3; *, p < 0.05 and **, p < 0.01). G. Measurement of ATP content in WT, Nrf1α-KO, and Nrf1α-OE cells following glutamine deprivation using a kit. The data are shown as fold changes (the mean ± SEM, n = 3 × 3; *, p < 0.05 and **, p < 0.01). H. Measurement of lactate levels in WT, Nrf1α-KO, and Nrf1α-OE cells following glutamine deprivation using a kit. The data are shown as fold changes (the mean ± SEM, n = 3 × 3; *, p < 0.05 and **, p < 0.01). I. Oil Red O staining of WT, Nrf1α-KO, and Nrf1α-OE cells following glutamine deprivation (with an original magnification of 200x). J. Metabolomic analysis of WT and Nrf1α-KO cells, with SMDP enrichment analysis results for differentially abundant metabolites compared to WT cells. K. Metabolomic analysis of WT and Nrf1α-KO cells, with KEGG pathway enrichment analysis results for differentially abundant metabolites compared to WT cells.

After glutamine deprivation, PKM, LDHA, and LDHD were slightly upregulated in all three cell types(Fig.4A and Fig. S2 C). Lactic acid and pyruvate, metabolites of glycolysis, were measured using a kit. Compared to WT cells, Nrf1α-KO cells exhibited significantly higher levels of lactic acid (Fig.4H) and pyruvate (Fig.4E), whereas Nrf1α-OE cells showed no significant change. Following glutamine deprivation, the levels of lactic acid (Fig.4H) and pyruvate (Fig.4E) decreased across all three cell lines.

The expression of key genes involved in the TCA cycle and oxidative phosphorylation (MT-OC1/2) was further investigated. RT-qPCR and Western blot analyses revealed that compared to WT cells, the expression of pyruvate dehydrogenase (PDHA1) increased in both Nrf1α-KO and Nrf1α-OE cells; however, glutamine deprivation did not affect PDHA1 expression. Citrate synthase (CS) expression remained unchanged across all conditions. Isocitrate dehydrogenase (IDH1/2 and IDH3B) expression decreased in Nrf1α-KO cells but increased in Nrf1α-OE cells. Following glutamine deprivation, IDH1 and IDH3B levels in WT cells remained stable, while IDH2 protein content increased. In Nrf1α-KO cells, the levels of IDH1, IDH2, and IDH3B were reduced. In Nrf1α-OE cells, IDH1 expression decreased, IDH2 increased, and IDH3B remained unchanged. Compared to WT cells, malate dehydrogenase (MDH1) and MT-OC1 gene expression did not significantly change in Nrf1α-KO or Nrf1α-OE cells, whereas MT-OC2 expression decreased in Nrf1α-KO cells but not in Nrf1α-OE cells. Notably, after glutamine deprivation, MDH1 and MT-OC1/2 expression levels significantly decreased in Nrf1α-KO cells, while MT-OC2 expression slightly decreased in WT and Nrf1α-OE cells. However, MDH1 and MT-OC1 expression levels in Nrf1α-OE and WT cells were unaffected by glutamine deprivation(Fig.4D and Fig.S2 D-E). Intracellular ATP levels were measured using a commercial kit. Results showed that compared to WT controls, ATP content increased in Nrf1α-KO and Nrf1α-OE cells. After glutamine starvation, ATP levels in WT and Nrf1α-OE cells remained stable, while ATP content in Nrf1α-KO cells significantly decreased(Fig.4G).These findings suggest that Nrf1α knockout leads to enhanced glycolysis and increased mitochondrial respiratory pressure, affecting mitochondrial number, morphology, and membrane potential following glutamine deprivation.

RT-qPCR and WB results demonstrated that, compared with the WT group, key lipid metabolism genes including ATP citrate lyase (ACLY), acetyl-CoA carboxylase (ACCA), fatty acid synthase (FASN), sterol regulatory element-binding proteins (SREBP1/2), members of the acyl-CoA synthetase long-chain family (ACSL3/4), stearoyl-CoA desaturase-1 (SCD1), and carnitine palmitoyltransferase (CPT) were significantly upregulated in Nrf1α-KO cells. However, in Nrf1α-OE cells, no significant changes were observed for most of these genes except for an increase in ACSL4 expression and a decrease in CPT expression. Following glutamine deprivation, ACCA expression decreased in all three cell types. In Nrf1α-KO cells, SREBP1/2, ACSL3, CPT, and SCD1 gene expression also decreased, while other genes remained unchanged (Fig.4C and Fig.S3A-B). To further investigate lipid metabolism, we conducted Oil Red O staining and triglyceride (TG) content detection. The TG content was highest in Nrf1α-KO cells, followed by Nrf1α-OE cells, with the lowest levels observed in the WT group. After glutamine deprivation, TG content in Nrf1α-KO cells significantly decreased, while itslightly decreased in the WT group and remained unchanged in Nrf1α-OE cells(Fig.4F). Oil Red O staining revealed substantial lipid accumulation in Nrf1α-KO cells, which was alleviated following glutamine deprivation(Fig.4I). In conclusion, Nrf1α knockout promotes lipid synthesis in Nrf1α-KO cells, whereas glutamine deprivation impairs this process. Nrf1α plays a crucial role in linking glutamine metabolism to lipid metabolism. Metabolomic analysis of Nrf1α-KO cells revealed 91 significantly altered metabolites compared to the WT group, comprising 36 upregulated and 55 downregulated metabolites((Fig.S3 E). These differentially expressed metabolites were enriched in pathways related to glutamate metabolism (SMDP) and lipid metabolism(Fig.4J), particularly lipid synthesis (KEGG)(Fig.4K). This evidence further supports the involvement of Nrf1α in regulating both lipid and glutamine metabolism.

Acetyl-CoA (AcCoA) serves as a central biosynthetic precursor for fatty acid synthesis and protein acetylation[19]. In cells, Glutamine is transported into mitochondria via the SLC1A5 transplasmic membrane transporter[20].glutamine is converted to glutamate in the mitochondria by glutaminase (GLS), an amide hydrolase that catalyzes this conversion while releasing ammonium ions[21]. The mitochondrial glutamate generated through these catabolic pathways can be exported from the mitochondria to the cytosol via the SLC25A18 and SLC25A22 transporters[20]; Once in the cytosol, glutamate contributes to glutathione biosynthesis and serves as an exchange factor for the import of extracellular cystine via SLC7A11(Fig.S1). Additionally, mitochondrial glutamate is converted to alpha-ketoglutarate (α-KG) by glutamate dehydrogenase 1 (GLUD1) or several mitochondrial aminotransferases, including alanine aminotransferase 2 (GPT2) and aspartate aminotransferase 2 (GOT2) (Fig.S1). α-KG is then transported from the mitochondria to the cytosol via SLC25A11, where it participates in fatty acid biosynthesis and NADH production[20, 22]. Mitochondrial α-KG can also enter the TCA cycle, supporting either the oxidative phosphorylation (OXPHOS) pathway or the reductive carboxylation pathway[23]. In the TCA cycle, AKG is converted to succinyl-CoA by ketoglutarate dehydrogenase (α-KGDH). Importantly, there is no net production of AKG in this cycle, and the AKG generated cannot be utilized for amino acid biosynthesis or other cellular functions. Research has demonstrated that tumor cells with mitochondrial dysfunction predominantly use glutamine-dependent reductive carboxylation instead of oxidative metabolism as the primary pathway for citrate synthesis. Specifically, AKG is reductively metabolized to citrate via IDH1 [24]. In this study, we examined the expression levels of genes related to glutamine metabolism using Western Blot (WB) and RT-qPCR. Experimental results showed that the expression level of the glutamine transporter ASCT2 was consistent across three cell lines and decreased in all after glutamine deprivation. There were no significant changes in the expression levels of the glutaminolytic gene GLUD; however, there was a significant increase in GLS and GPT in Nrf1α knockout (Nrf1α-KO) cells, with a slight increase observed in Nrf1α overexpressing (Nrf1α-OE) cells. After glutamine deprivation, GLS levels significantly decreased in all three cell lines, while GPT slightly decreased only in Nrf1α-KO cells, with no significant change in WT and Nrf1α-OE cells. Additionally, the expression levels of other amino acid metabolic genes PSAT, ASS1, PHGDH were significantly increased in Nrf1α-KO cells. The expression of the cystine/glutamate reverse transporter xCT was elevated in both Nrf1α-KO and Nrf1α -OE cells. In wild-type (WT) cells, xCT expression was significantly increased following glutamine deprivation; however, no significant changes were observed in Nrf1α-KO and Nrf1α-OE cells under the same conditions. This indicates that knocking out the Nrf1α gene promotes the consumption of glutamine. Interestingly, compared to wild-type (WT) cells, the expression level of the glutamine synthetase gene GLUL was significantly lower in Nrf1α-KO cells but significantly higher in Nrf1α-OE cells. After glutamine deprivation, the expression level of GLUL was highest in Nrf1α-OE cells, followed by WT cells, and lowest in Nrf1α-KO cells(Fig.4D and Fig.S3C-D). In summary, we hypothesized that Nrf1α gene knockout leads to two major metabolic alterations. First, the enhancement of glycolysis imposes additional respiratory pressure on mitochondria. Second, the upregulation of lipid synthesis results in a significant reduction and metabolism of α-KG by IDH1 into citrate, which is transported from mitochondria to the cytoplasm for lipid synthesis, further exacerbating mitochondrial respiratory pressure. Consequently, cells must activate the glutamine catabolism pathway to replenish mitochondrial α-KG levels, leading to substantial glutamine decomposition and consumption. Additionally, Nrf1α gene knockout decreases GLUL expression, reducing cellular glutamine synthesis. Collectively, these factors result in extensive utilization of α-KG for lipid metabolism upon glutamine deprivation, thereby disrupting the mitochondrial TCA cycle, impairing oxidative phosphorylation, and decreasing intracellular GSH content, ultimately causing redox imbalance and apoptosis.

### 3.5 The Nrf1α gene regulates apoptosis after glutamine deprivation by influencing the expression of the GLUL gene

Nrf1 functions as a transcription factor that binds to antioxidant response elements (AREs) and/or electrophilic response elements (EpREs) within the promoter regions of target genes, thereby modulating the expression of downstream target genes[25]. Through the analysis of the GLUL promoter region sequence, we discovered an ARE (Antioxidant Response Element) site within the GLUL promoter region. The experiment involved cloning the GLUL promoter region into the PGL3-Basic vector for a dual luciferase reporter assay, which showed that Nrf1α can activate the transcription of the GLUL gene(Fig.5A). Immunocytochemical confocal imaging revealed that compared to the wild-type (WT) group, Nrf1α protein expression was reduced along with GLUL protein expression in Nrf1α knockout (Nrf1α-KO) cells, while Nrf1α protein expression increased alongside GLUL protein expression in Nrf1α overexpressing (Nrf1α-OE) cells(Fig.5B). This indicates that Nrf1α promotes the expression of the glutamine synthesis gene GLUL. To further investigate this, we constructed the coding sequence (CDS) of GLUL into the gene expression vector pcDNA3.1 to create a GLUL overexpression vector and transfected it into Nrf1 α-KO cells. Western blot (WB) results confirmed successful transfection, showing that Nrf1α-KO cells could overexpress the GLUL protein(Fig.5C). We then subjected these GLUL-overexpressing Nrf1α-KO cells to glutamine starvation. Experimental results demonstrated that after 24 hours of glutamine deprivation, ROS levels were lower in cells overexpressing GLUL compared to control cells (transfected with empty pcDNA3.1 vector) (Fig.5E), but GSH levels did not show significant changes, nor did cell apoptosis rates(Fig.5D and 5G). Interestingly, after 24 hours of glutamine deprivation, the mitochondrial membrane potential in GLUL-overexpressing Nrf1α-KO cells was higher than in the control group(Fig.5F). When glutamine deprivation was extended to 48 hours, a large number of control cells (transfected with empty pcDNA3.1 vector) died, whereas GLUL expression effectively alleviated cell death in Nrf1α-KO cells(Fig.5H). WB experiments further showed that after 24 hours of glutamine deprivation, the expression levels of apoptosis-related genes Caspase-9 and Caspase-3 were lower in Nrf1α-KO cells expressing GLUL compared to the control group, while Bax and BCL2 levels did not show significant changes(Fig.5I).

**Fig. 5.**
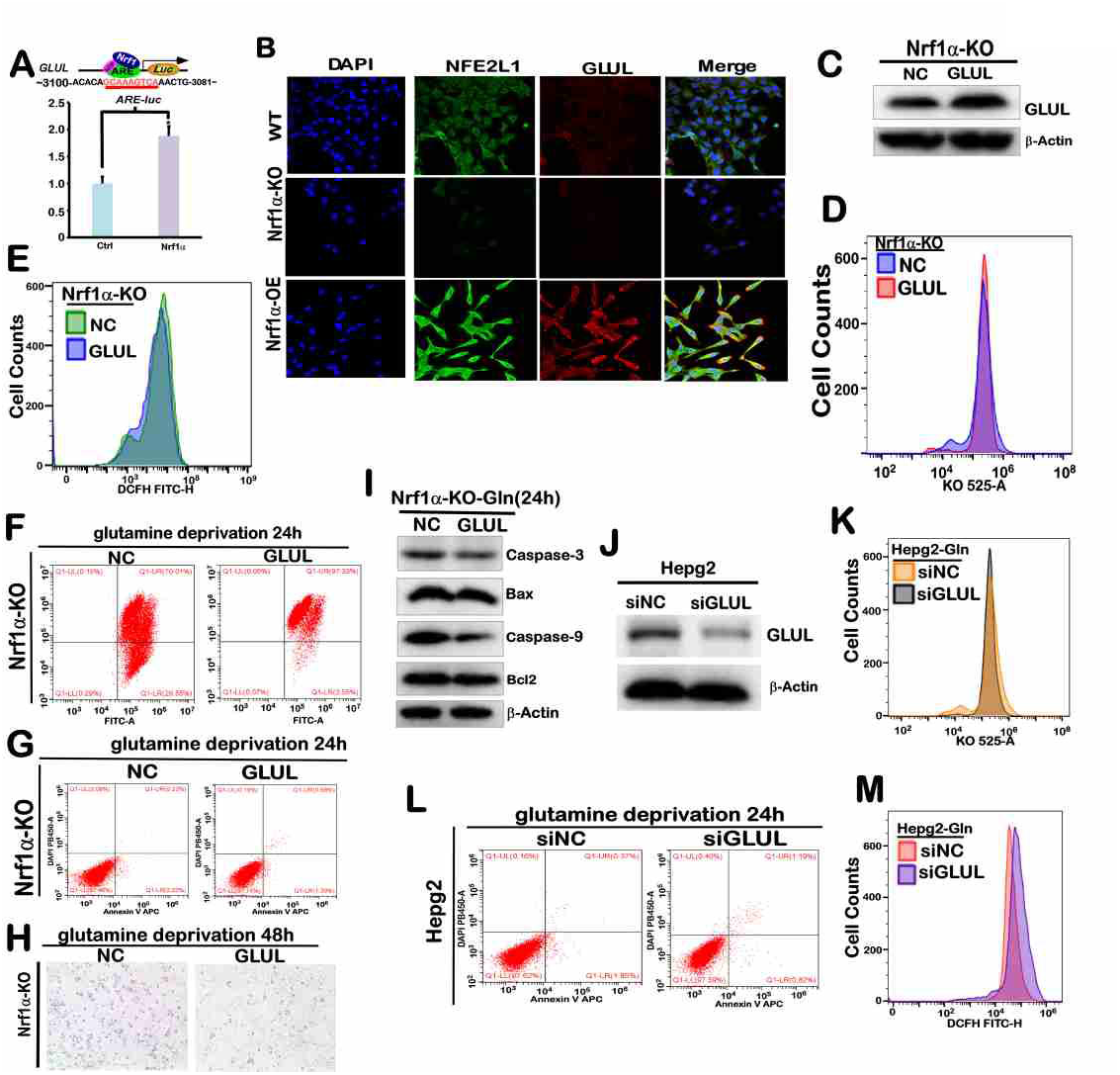
The Nrf1α gene regulates apoptosis after glutamine deprivation by influencing the expression of the GLUL gene. A. Cells were transfected with the GLUL-pGL3 promoter vector and pRL-TK along with Nrf1 expression constructs (or empty pcDNA3.1 plasmid), and after a 24-hour recovery, luciferase activity was measured using a dual-luciferase reporter assay kit. The data are shown as fold changes (the mean ± SEM, n = 3 × 3; *, p < 0.05 and **, p < 0.01). B. Immunocytochemistry was performed on WT, Nrf1α-KO, and Nrf1α-OE cells using primary antibodies against Nrf1α and GLUL and fluorescent secondary antibodies, followed by DAPI staining. Fluorescence images were captured and merged (scale bar = 20 μm). C. Western blotting was used to detect GLUL protein expression in Nrf1α-KO cells transfected with the GLUL expression plasmid after a 24-hour recovery period and subsequent 24 hours of glutamine starvation. D. Changes in cellular GSH levels were detected by flow cytometry in Nrf1α-KO cells transfected with the GLUL expression plasmid, following a 24-hour recovery period, 24 hours of glutamine starvation, and 30 minutes of mBCL staining. E. Changes in cellular ROS levels were detected by flow cytometry in Nrf1α-KO cells transfected with the GLUL expression plasmid, following a 24-hour recovery period, 24 hours of glutamine starvation, and 30 minutes of DCFH staining. F. Mitochondrial membrane potential was analyzed by flow cytometry in Nrf1α-KO cells transfected with the GLUL expression plasmid, following a 24-hour recovery period, 24 hours of glutamine starvation, and 30 minutes of JC-1 incubation. G. Cell apoptosis was analyzed by flow cytometry in Nrf1α-KO cells transfected with the GLUL expression plasmid, following a 24-hour recovery period, 24 hours of glutamine starvation, and incubation with Annexin V-FITC and PI. H. Cellular morphology was observed under a microscope in Nrf1α-KO cells transfected with the GLUL expression plasmid after a 24-hour recovery period and 48 hours of glutamine starvation (with an original magnification of 200x). I. Expression levels of Caspase-3, Caspase-9, Bax, and BCL2 proteins were detected by Western blotting in Nrf1α-KO cells transfected with the GLUL expression plasmid after a 24-hour recovery period and 24 hours of glutamine starvation. J. GLUL protein expression was detected by Western blotting in Nrf1α-KO cells transfected with siRNA targeting GLUL, after a 24-hour recovery period and 24 hours of glutamine starvation. K. Changes in cellular GSH levels were detected by flow cytometry in Nrf1α-KO cells transfected with siRNA targeting GLUL, after a 24-hour recovery period, 24 hours of glutamine starvation, and 30 minutes of mBCL staining. L. Cell apoptosis was analyzed by flow cytometry in Nrf1α-KO cells transfected with siRNA targeting GLUL, after a 24-hour recovery period, 24 hours of glutamine starvation, and incubation with Annexin V-FITC and PI. M. Changes in cellular ROS levels were detected by flow cytometry in Nrf1α-KO cells transfected with siRNA targeting GLUL, after a 24-hour recovery period, 24 hours of glutamine starvation, and 30 minutes of DCFH staining.

Additionally, we used specific siRNA (F:5’-GAUUGGACCUUGUGAAGGAdTdT-3’, R:5’ -UCCUUCACAAGCUCCAAUCdTdT-3’) to knock down the expression of GLUL protein in wild-type (WT) cells. Western blot (WB) results showed that GLUL protein expression was reduced in the siGLUL group compared to the siNC (negative control) group. We then subjected both the siGLUL and siNC groups to glutamine starvation. Experimental results indicated that after 24 hours of glutamine deprivation, the ROS levels were higher in the siGLUL group compared to the siNC group(Fig.5M), while GSH content did not show significant changes(Fig.5K), nor did cell apoptosis rates(Fig.5L).The above experimental results indicate that the Nrf1 α gene can regulate the expression of the glutamine synthetase GLUL gene. Therefore, one of the reasons for the significant cell death observed in Nrf1α knockout (Nrf1α-KO) cells after glutamine deprivation is the reduced expression of the glutamine synthetase gene GLUL.

### 3.6 Supplementation with a-KG can mitigate the rapid apoptosis of Nrf1α-KO cells caused by glutamine deprivation

The metabolite alpha-ketoglutaric acid (α-KG) is a key intermediate in the TCA cycle and plays a crucial role in various metabolic and cellular pathways[15]. To investigate whether mitochondrial dysfunction in Nrf1α-KO cells results in intensified oxidative stress and rapid apoptosis following glutamine deprivation, we hypothesized that this phenomenon may be attributed to the substantial influx of α-KG into the cytoplasm, which subsequently participates in lipid synthesis metabolism. We conducted experiments to evaluate the effects of exogenous α-KG supplementation. Since exogenously added α-KG cannot cross the cell membrane, we utilized dimethyl alpha-ketoglutarate (DM-αKG), a cell-permeable analog of α-KG, to supplement intracellular α-KG levels[26]. The results demonstrated that exogenous supplementation of DM-αKG (15 mM) alleviated amino acid deprivation-induced apoptosis in Nrf1α-KO cells and increased mitochondrial membrane potentia (l Fig.6B-C). Under glutamine deprivation, a significant increase in cell death was observed in Nrf1α-KO cells after 48 hours of microscopy examination. However, glutamine deprivation combined with DM-αKG (15 mM) supplementation did not result in significant cell death (Fig.6A). Further experiments revealed that DM-αKG supplementation decreased ROS levels (Fig.6D), increased TG content(Fig.6E), and elevated ATP levels in Nrf1α-KO cells(Fig.6G), while GSH levels remained unchanged(Fig.6F). Western blot analysis showed reduced expression of pro-apoptotic proteins Caspase-9, Caspase-3, and Bax, and increased expression of the anti-apoptotic protein BCL2 in Nrf1α-KO cells supplemented with DM-αKG during glutamine deprivation(Fig.6H). No significant changes were observed in the expression of lipid metabolism-related genes SREBP1/2, FASN, ACLY, ACACA, CD36, and ACSL3/4(Fig.6I). Protein expression of mitochondrial metabolism-related genes IDH1, IDH3B, SDHA, SDHB, MDH1, MT-OC1, and MT-OC2 increased, although IDH2 expression did not change significantly(Fig.6J). The aforementioned experimental findings suggest that following the knockout of the Nrf1α gene, a significant quantity of glutamine is catabolized into α -ketoglutarate (α -KG) to sustain mitochondrial metabolism. Consequently, upon glutamine deprivation, the intramitochondrial α-KG levels in cells decline precipitously, becoming inadequate to support mitochondrial metabolic processes. Additionally, the glutamine content in Nrf1α-KO cells diminishes markedly, insufficient to support cellular glutathione (GSH) synthesis, leading to an elevation in cellular reactive oxygen species (ROS). Ultimately, this culminates in apoptosis.

**Fig. 6.**
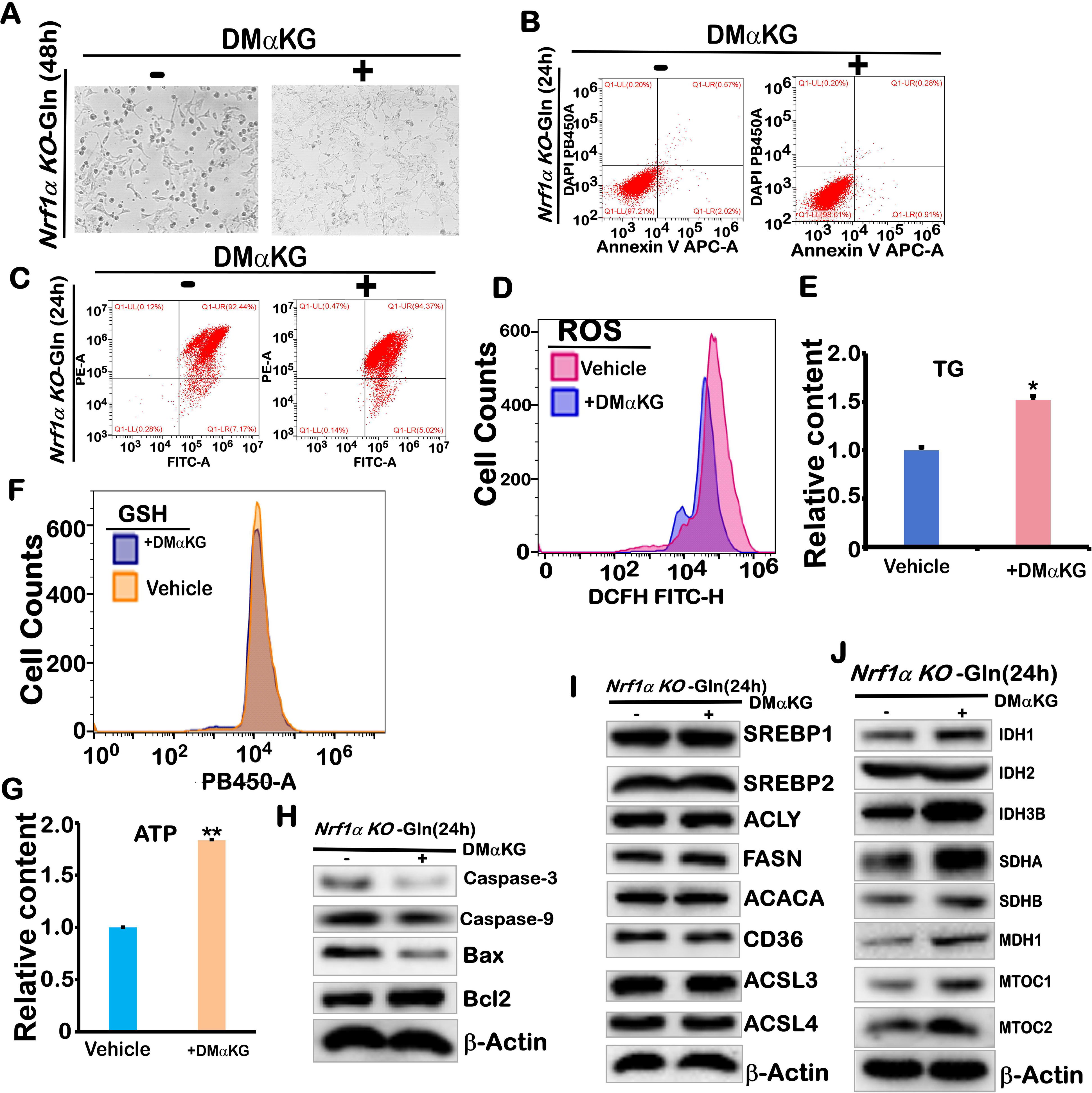
Supplementation with α-KG alleviates rapid apoptosis in Nrf1α-KO cells induced by glutamine deprivation. A. Morphology of Nrf1α-KO cells cultured for 48 hours in glutamine-free medium containing DM-αKG (15 mM) was observed under a microscope (with an original magnification of 200x). B. Apoptosis analysis of Nrf1α-KO cells cultured for 24 hours in glutamine-free medium containing DM-αKG (15 mM) using flow cytometry after incubation with Annexin V-FITC and PI. C. Mitochondrial membrane potential analysis of Nrf1α-KO cells cultured for 24 hours in glutamine-free medium containing DM-αKG (15 mM) using flow cytometry after 30 minutes of JC-1 incubation. D. Changes in cellular ROS levels were detected by flow cytometry in Nrf1α-KO cells cultured for 24 hours in glutamine-free medium containing DM-αKG (15 mM), following 30 minutes of DCFH staining. E. Triglyceride (TG) content in Nrf1α-KO cells cultured for 24 hours in glutamine-free medium containing DM-αKG (15 mM) was measured using a kit. The data are shown as fold changes (the mean ± SEM, n = 3 × 3; *, p < 0.05 and **, p < 0.01). F. Changes in cellular GSH levels were detected by flow cytometry in Nrf1α-KO cells cultured for 24 hours in glutamine-free medium containing DM-αKG (15 mM), following 30 minutes of mBCL staining. G. ATP content in Nrf1α-KO cells cultured for 24 hours in glutamine-free medium containing DM-αKG (15 mM) was measured using a kit. The data are shown as fold changes (the mean ± SEM, n = 3 × 3; *, p < 0.05 and **, p < 0.01). H. Expression levels of Caspase-3, Caspase-9, Bax, and BCL2 proteins were detected by Western blotting in Nrf1α-KO cells cultured for 24 hours in glutamine-free medium containing DM-αKG (15 mM). I. Expression levels of SREBP1, SREBP2, ACLY, FASN, ACACA, CD36, ACSL3, and ACSL4 proteins were detected by Western blotting in Nrf1α-KO cells cultured for 24 hours in glutamine-free medium containing DM-αKG (15 mM). J. Expression levels of IDH1, IDH2, IDH3B, SDHA, SDHB, MDH1, MTOC1, and MTOC2 proteins were detected by Western blotting in Nrf1α-KO cells cultured for 24 hours in glutamine-free medium containing DM-αKG (15 mM).”

### 3.7 Inhibition of fatty acid synthesis can mitigate the rapid apoptosis of Nrf1α-KO cells induced by glutamine deprivation

To determine whether the massive death of Nrf1α-KO cells under glutamine deprivation was caused by the extensive catabolism of glutamine into α-ketoglutarate (α-KG) for fatty acid synthesis, leading to insufficient intracellular GSH synthesis and increased ROS levels that induce apoptosis, we utilized Fatostatin[27], an SREBP inhibitor, and C75[28], a FASN inhibitor, to inhibit the fatty acid synthesis pathway in glutamine-deprived Nrf1α-KO cells. WB experiments demonstrated that C75 significantly inhibited the expression of fatty acid synthetase (FASN) protein in wild-type (WT) and NRF1-α knockout (KO) cells under glutamine deprivation (Fig.7L). Apoptosis detection results indicated that inhibiting FASN protein expression for 24 hours under glutamine deprivation effectively alleviated Nrf1α-KO cell apoptosis (Fig.7A). Extending the time to 48 hours revealed substantial cell death in the DMSO control group, whereas the C75 group significantly mitigated cell death (Fig.7C). Additionally, inhibition of FASN expression by glutamine deprivation reduced triglyceride (TG) content (Fig.7F), increased mitochondrial membrane potential(Fig.7H), decreased reactive oxygen species (ROS) (Fig.7G), and increased glutathione (GSH) (Fig.7K) levels in NRF1-α KO cells. WB analysis showed that Fatostatin treatment decreased the protein expressions of SREBP1, SREBP2, ACLY, FASN, and ACACA in glutamine-deprived WT and NRF1-α KO cells(Fig.7M), leading to a reduction in TG content in NRF1-α KO cells (Fig.7E). These findings suggest that Fatostatin can inhibit lipid synthesis. Furthermore, Fatostatin treatment in glutamine-deprived NRF1-α KO cells reduced apoptosis(Fig.7B), increased mitochondrial membrane potential(Fig.7H), decreased ROS(Fig.7J), and increased GSH content(Fig.7I). Western blotting detected changes in the expression of apoptosis-related genes in glutamine-deprived WT and NRF1-α KO cells treated with C75 and Fatostatin inhibitors. The results showed that C75 inhibited the expression of pro-apoptotic proteins Caspase-9, Caspase-3, and Bax in both WT and NRF1-α KO cells after lipid metabolism inhibition, while increasing anti-apoptotic BCL2 expression(Fig.7L). In contrast, Fatostatin treatment did not significantly alter the expression of these proteins in WT cells but decreased the expression of Caspase-9, Caspase-3, and Bax and increased BCL2 expression in glutamine-deprived NRF1-α KO cells (Fig.7N). The experimental results demonstrated that inhibiting lipid metabolism effectively restored the mitochondrial membrane potential decline induced by amine deficiency in NRF1-α-KO cells. Moreover, this inhibition significantly mitigated the accumulation of reactive oxygen species (ROS) and slowed the apoptosis process in NRF1-α-KO cells under glutamine-deficient conditions. Further investigations revealed that following NRF1-α gene knockout, a substantial amount of glutamine is redirected into the mitochondria for α-ketoglutarate (α-KG) synthesis. Consequently, insufficient glutamine supply leads to rapid depletion of mitochondrial α-KG, severely impacting mitochondrial metabolic function. This impairment prevents adequate glutathione (GSH) synthesis, resulting in elevated ROS levels and ultimately inducing apoptosis.

**Fig. 7.**
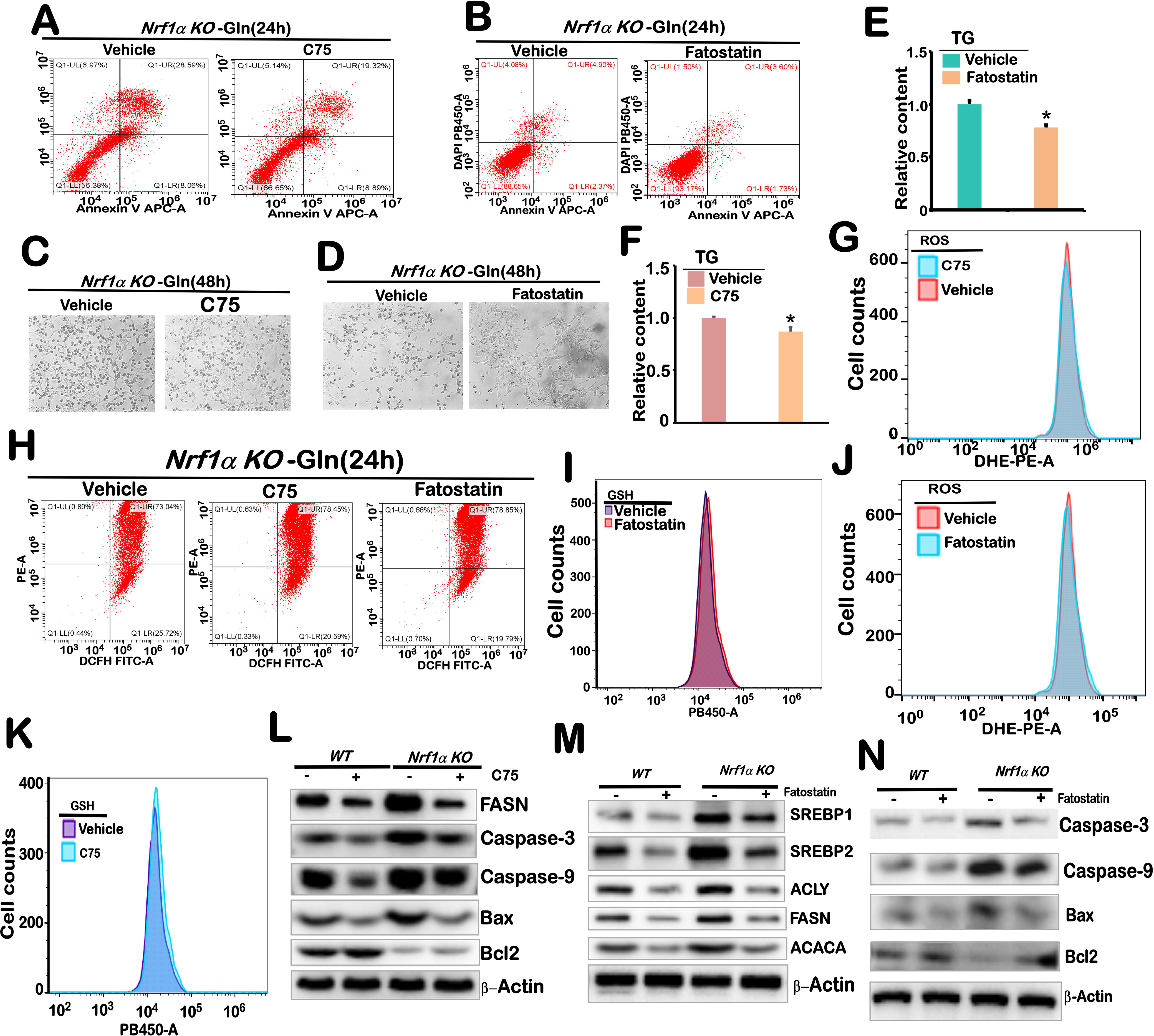
Inhibition of lipid metabolism can alleviate rapid apoptosis in Nrf1α-KO cells induced by glutamine deprivation. A. Apoptosis analysis of Nrf1α-KO cells cultured for 24 hours in glutamine-free medium containing C75 (10 μM) using flow cytometry after incubation with Annexin V-FITC and PI. B. Apoptosis analysis of Nrf1α-KO cells cultured for 24 hours in glutamine-free medium containing fatostatin (20 μM) using flow cytometry after incubation with Annexin V-FITC and PI. C. Morphology of Nrf1α-KO cells cultured for 48 hours in glutamine-free medium containing C75 (10 μM) was observed under a microscope (with an original magnification of 200x). D. Morphology of Nrf1α-KO cells cultured for 48 hours in glutamine-free medium containing fatostatin (20 μM) was observed under a microscope (with an original magnification of 200x). E. Triglyceride (TG) content in Nrf1α-KO cells cultured for 24 hours in glutamine-free medium containing fatostatin (20 μM) was measured using a kit. The data are shown as fold changes (the mean ± SEM, n = 3 × 3; *, p < 0.05 and **, p < 0.01). F. Triglyceride (TG) content in Nrf1α-KO cells cultured for 24 hours in glutamine-free medium containing C75 (10 μM) was measured using a kit. The data are shown as fold changes (the mean ± SEM, n = 3 × 3; *, p < 0.05 and **, p < 0.01). G. Changes in cellular ROS levels were detected by flow cytometry in Nrf1α-KO cells cultured for 24 hours in glutamine-free medium containing C75 (10 μM), following 30 minutes of DHE staining. H. Mitochondrial membrane potential analysis of Nrf1α-KO cells cultured for 24 hours in glutamine-free medium containing C75 (10 μM) or fatostatin (20 μM) using flow cytometry after 30 minutes of JC-1 incubation. I. Changes in cellular GSH levels were detected by flow cytometry in Nrf1α-KO cells cultured for 24 hours in glutamine-free medium containing fatostatin (20 μM), following 30 minutes of mBCL staining. J. Changes in cellular ROS levels were detected by flow cytometry in Nrf1α-KO cells cultured for 24 hours in glutamine-free medium containing fatostatin (20 μM), following 30 minutes of DHE staining. K. Changes in cellular GSH levels were detected by flow cytometry in Nrf1α-KO cells cultured for 24 hours in glutamine-free medium containing C75 (10 μM), following 30 minutes of mBCL staining. L. Expression levels of FASN, Caspase-3, Caspase-9, Bax, and BCL2 proteins were detected by Western blotting in WT and Nrf1α-KO cells cultured for 24 hours in glutamine-free medium containing C75 (10 μM). M. Expression levels of SREBP1, SREBP2, ACLY, FASN, and ACACA proteins were detected by Western blotting in WT and Nrf1α-KO cells cultured for 24 hours in glutamine-free medium containing fatostatin (20 μM). N. Expression levels of Caspase-3, Caspase-9, Bax, and Bcl-2 proteins were detected by Western blotting in WT and Nrf1α-KO cells cultured for 24 hours in glutamine-free medium containing fatostatin (20 μM).

### 3.8 Inhibition of the PI3K-AKT-mTOR signaling pathway can mitigate apoptosis induced by glutamine deprivation in Nrf1α-KO cells

The PI3K/Akt/mTOR signaling pathway is a critical intracellular signal transduction pathway that enhances lipid synthesis via multiple mechanisms[29, 30]. WB and RT-qPCR results demonstrated that, compared with WT cells, the expression levels of PI3K, p-AKT, p-mTOR, and AKT were elevated in Nrf1α-KO cells. Conversely, only the expression of p-mTOR was increased in Nrf1α-OE cells, while no significant changes ere observed for PI3K, p-AKT, and AKT(Fig.8A). To investigate the effects of inhibiting the PI3K-AKT-mTOR signaling pathway in glutamine-deprived Nrf1α-KO cells, we used NVP-BKM120 (NVP) and rapamycin (RAPA). WB analysis revealed that treatment with NVP under glutamine deprivation conditions led to decreased expression of p-mTOR and p-AKT proteins, whereas PI3K protein levels remained unchanged(Fig.8C). When combining glutamine deprivation with RAPA treatment, the expression of p-mTOR protein was reduced in Nrf1α-KO cells, while p-AKT and PI3K levels did not change significantly(Fig.8C). The results indicated that both NVP and RAPA inhibitors effectively suppressed the expression of SREBP1, SREBP2, ACLY, and FASN proteins involved in lipid synthesis(Fig.8H). Both inhibitors also alleviated apoptosis induced by glutamine deprivation in Nrf1α-KO cells(Fig.8B). Microscopic observations after 48 hours of treatment with NVP and RAPA inhibitors showed a reduction in cell death(Fig.8D). Further experiments revealed no significant changes in GSH levels in glutamine-deprived Nrf1α-KO cells treated with NVP(Fig.8F), but ROS levels were slightly decreased(Fig.8I). In contrast, treatment with RAPA inhibitors did not significantly alter GSH (Fig.8G) levels but significantly reduced ROS levels(Fig.8J). This may be attributed to the utilization of intracellular GSH for ROS reduction. Experimental results confirmed that NVP and RAPA inhibitors significantly reduced intracellular TG content in glutamine-deprived Nrf1α-KO cells(Fig.8E). These findings indicate that inhibiting the PI3K-AKT-mTOR signaling pathway can suppress fat synthesis, consequently preventing the extensive influx of glutamine into the mitochondria for degradation into α-ketoglutarate (αKG), which is utilized for the reverse synthesis of citric acid. This alleviates the excessive consumption of glutamine to replenish the citric acid cycle, further promoting glutathione synthesis from glutamine to mitigate oxidative stress in NRF1-αKO cells, and ultimately reducing cell apoptosis.

**Fig. 8.**
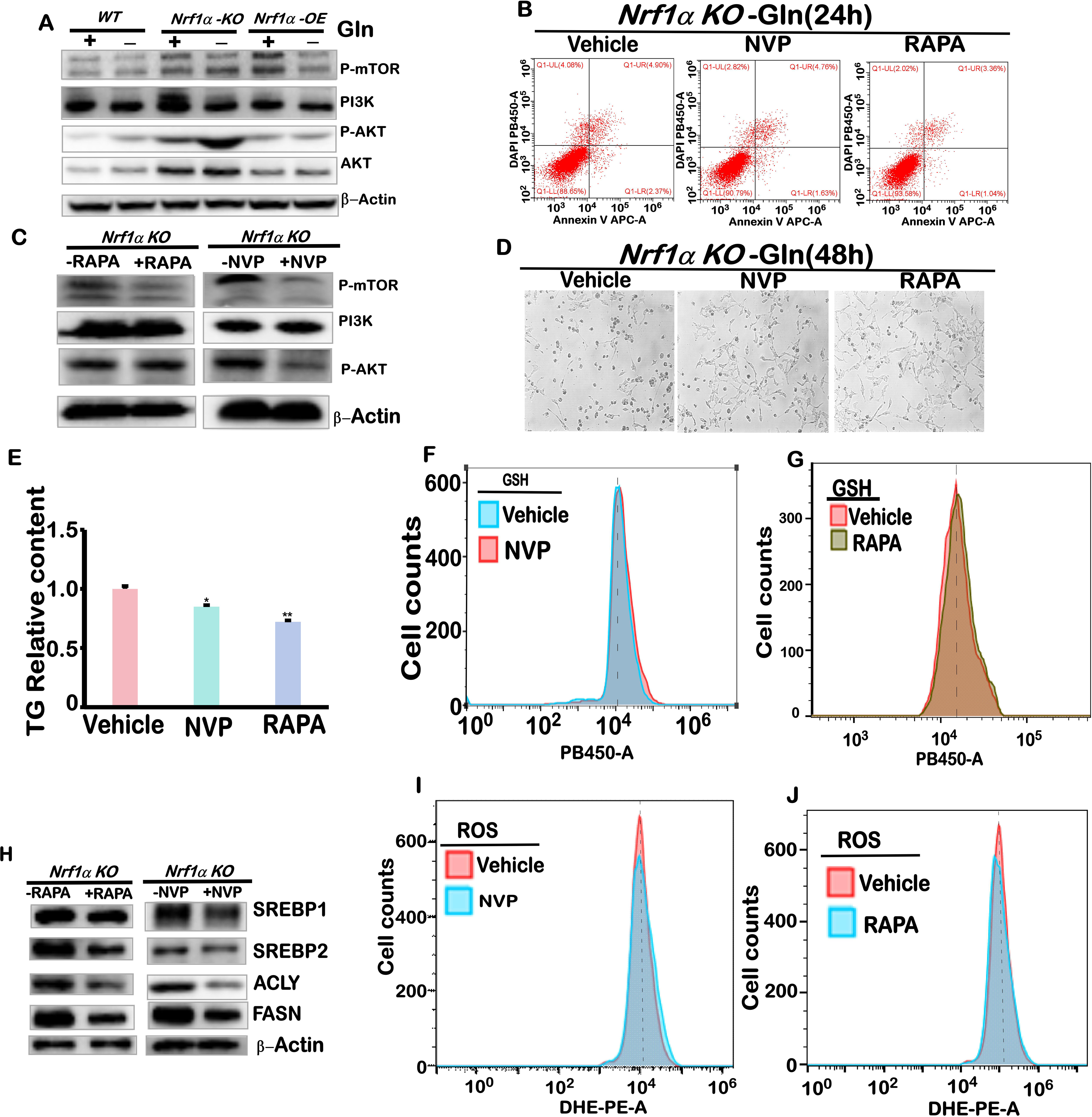
Inhibition of the PI3K-AKT-mTOR signaling pathway that regulates lipid metabolism can alleviate apoptosis in Nrf1α-KO cells caused by glutamine deprivation. A. Detection of PI3K, AKT, p-AKT, and p-mTOR protein expression in WT, Nrf1α-KO, and Nrf1α-OE cells after 24 hours of glutamine deprivation using Western blotting. B. Apoptosis analysis of Nrf1α-KO cells cultured for 24 hours in glutamine-free medium containing NVP-BKM120 (0.5 μM) or RAPA (200 nM) using flow cytometry after incubation with Annexin V-FITC and PI. C. Detection of PI3K, p-AKT, and p-mTOR protein expression in Nrf1α-KO cells cultured for 24 hours in glutamine-free medium containing NVP-BKM120 (0.5 μM) or RAPA (200 nM) using Western blotting. D. Morphology of Nrf1α-KO cells cultured for 48 hours in glutamine-free medium containing NVP-BKM120 (0.5 μM) or RAPA (200 nM) was observed under a microscope (with an original magnification of 200x). E. Triglyceride (TG) content in Nrf1α-KO cells cultured for 24 hours in glutamine-free medium containing NVP-BKM120 (0.5 μM) or RAPA (200 nM) was measured using a kit. F. Changes in cellular GSH levels were detected by flow cytometry in Nrf1α-KO cells cultured for 24 hours in glutamine-free medium containing NVP-BKM120 (0.5 μM), following 30 minutes of mBCL staining. G. Changes in cellular GSH levels were detected by flow cytometry in Nrf1α-KO cells cultured for 24 hours in glutamine-free medium containing RAPA (200 nM), following 30 minutes of mBCL staining. H. Expression levels of SREBP1, SREBP2, ACLY, and FASN proteins were detected by Western blotting in WT and Nrf1α-KO cells cultured for 24 hours in glutamine-free medium containing NVP-BKM120 (0.5 μM) or RAPA (200 nM). I. Changes in cellular ROS levels were detected by flow cytometry in Nrf1α-KO cells cultured for 24 hours in glutamine-free medium containing NVP-BKM120 (0.5 μM), following 30 minutes of DHE staining. J. Changes in cellular ROS levels were detected by flow cytometry in Nrf1α-KO cells cultured for 24 hours in glutamine-free medium containing RAPA (200 nM), following 30 minutes of DHE staining.

## 4. Discussion

Glutamine is an essential amino acid for cancer cells, with its consumption rate far exceeding the rate of biosynthesis[31]. It plays a critical role in replenishing the tricarboxylic acid (TCA) cycle, as well as in the biosynthesis of nucleotides, glutathione (GSH), and other non-essential amino acids within cellular metabolism. Glutamine starvation leads to the disruption of intracellular glutamine metabolic pathways, thereby affecting mitochondrial function and redox balance. For example, glutamine starvation reduces the intracellular concentration of glutamate derived from glutamine, leading to a decrease in mitochondrial membrane potential and an increase in reactive oxygen species (ROS) levels. This oxidative stress damages cellular components, such as through lipid peroxidation, ultimately triggering apoptosis. [32]. Our findings indicate that Nrf1α gene knockout in HepG2 cells enhances glycolysis, leading to increased intracellular glucose degradation and anaerobic glycolysis rather than mitochondrial aerobic oxidation. This shift results in weakened mitochondrial metabolism and reduced membrane potential. Additionally, Nrf1α gene knockout promotes fatty acid synthesis in HepG2 cells, causing substantial glutamine catabolism into α-ketoglutarate (αKG) within mitochondria. This process reverses the TCA cycle to produce citrate, which then enters the cytoplasm for lipid synthesis. These two factors lead to a reduction in mitochondrial αKG levels in a glutamine-deficient environment, further resulting in a sharp decline in αKG, disruption of the mitochondrial TCA cycle, and decreased mitochondrial membrane potential. Simultaneously, the depletion of glutamine lowers the intracellular GSH/ROS ratio, ultimately inducing apoptosis (Fig.9).

**Fig. 9.**
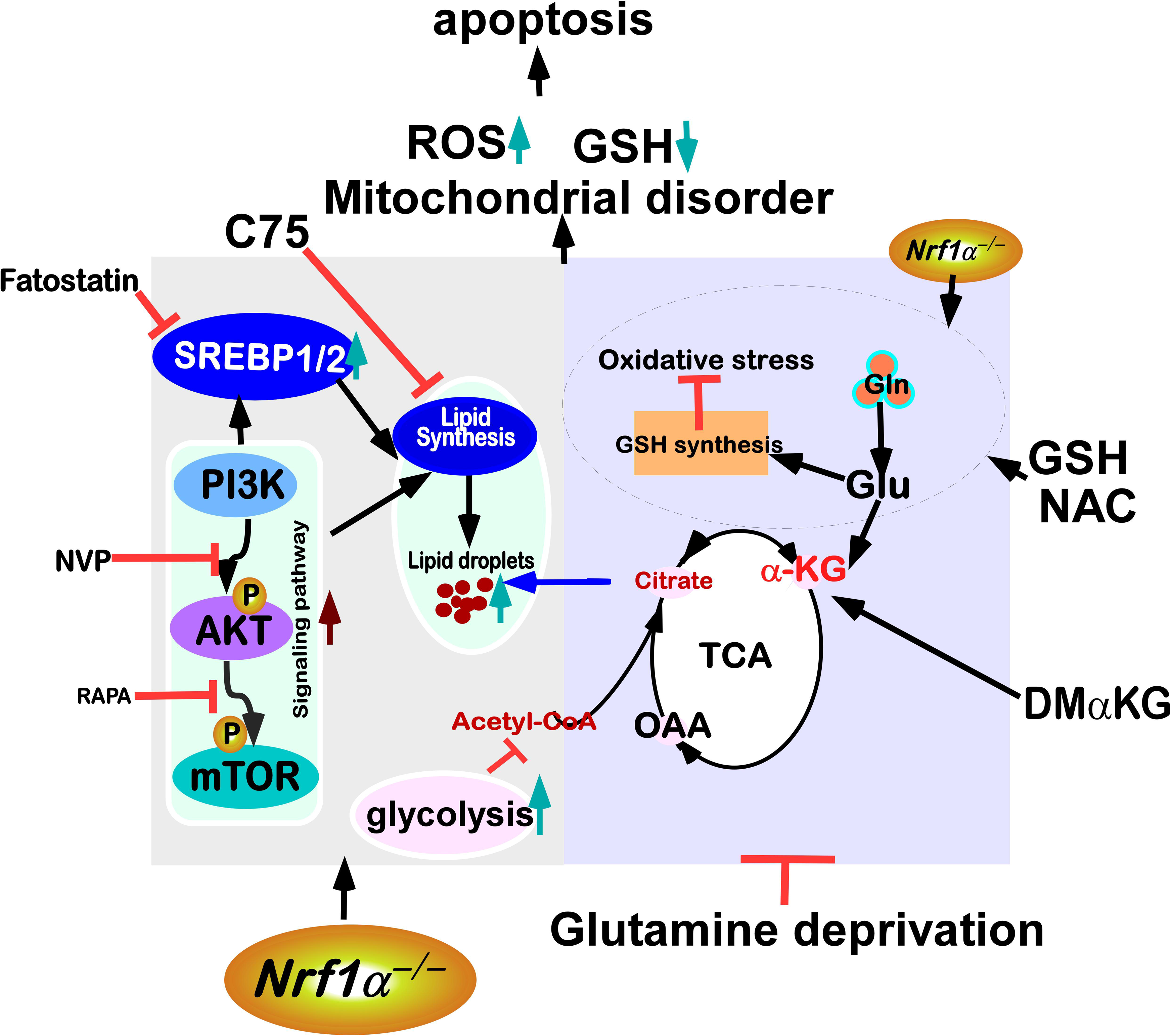
Genetic loss of Nrf1α promotes apoptosis of HepG2 cells during glutamine deprivation by triggering glutamine addiction.

Metabolic reprogramming has emerged as a hallmark of cancer[33], with the reconfiguration of glutamine metabolism playing a pivotal role, particularly in hepatocellular carcinoma (HCC)[34]. Extensive evidence demonstrates that glutamine deprivation can induce apoptosis in tumor cell lines by activating the intrinsic apoptotic pathway[35]. Our study revealed that the expression of Nrf1α protein in HepG2, Huh7, and MHCC97H cells initially increased and subsequently decreased. Glutamine deprivation resulted in rapid apoptosis in Nrf1α-knockout HepG2 cells. These findings indicate that Nrf1α is crucial for regulating glutamine metabolism (Fig.1A-C). Glutamine metabolism stimulates the biosynthesis of glutathione (GSH) and nicotinamide adenine dinucleotide phosphate (NADPH), thereby playing a crucial role in sustaining oxidative homeostasis in cancer cells[15]. Loss of Nrf1α not only results in severe oxidative stress but also enhances susceptibility to cytotoxicity induced by oxidative stress[36, 37]. In HepG2 cells with Nrf1α knockout, both GSH levels and ROS production were elevated. Notably, following glutamine starvation in these Nrf1α-deficient cells, ROS accumulation and intracellular GSH depletion were further exacerbated. However, supplementation with GSH and N-acetylcysteine (NAC) during glutamine starvation significantly reduced intracellular ROS levels (Fig.3). These findings indicate that Nrf1α plays a critical role in maintaining cellular redox homeostasis by modulating glutamine metabolism. Cells generate the maintenance bioenergy in the form of ATP through glycolysis and channel the remaining glucose-derived intermediates into the TCA cycle, where they are converted to citrate[38]. The TCA cycle serves as a central hub for energy metabolism, macromolecular synthesis, and redox balance. Nrf1-knockout HepG2 cells exhibit enhanced glycolysis, leading to a reduced flux of residual glucose into the TCA cycle. Lipid metabolism is the process by which cells convert nutrients into metabolic intermediates for use in membrane biosynthesis, energy storage, and the production of signaling molecules. Cancer cells meet their rapid proliferation demands through lipid metabolism, primarily acquiring lipids via two pathways: de novo synthesis: utilizing glucose or glutamine as substrates to generate fatty acids and cholesterol through synthetic pathways. Under hypoxic conditions, glutamine-derived α-ketoglutarate (α-KG) can be converted into citrate, promoting de novo fatty acid synthesis. Additionally, glutamine metabolism can support fatty acid synthesis by regulating NADPH generation, maintaining cellular redox balance. In Nrf1α-knockout HepG2 cells, citrate preferentially exits the mitochondria into the cytosol to support lipid synthesis. However, increased citrate efflux from the mitochondria consumes TCA cycle metabolites. Therefore, Nrf1α-knockout HepG2 cells enhance glutaminolysis to produce α-KG, which enters the TCA cycle to maintain mitochondrial TCA cycle activity and oxidative phosphorylation. (Fig4 and Fig6).

Glutamine is a major contributor to the anaerobic supplementation of the TCA cycle in rapidly growing tumor cells. This dependence on glutamine metabolism, commonly eferred to as glutamine addiction, plays multiple roles in cell proliferation by providing carbon and reductive nitrogen for biosynthetic reactions and redox homeostasis through the TCA cycle. The interplay between glutamine addiction and lipid metabolism offers new targets for cancer therapy. For example, inhibiting ASCT2 or key enzymes of glutamine metabolism (such as GLS) can reduce fatty acid synthesis, thereby suppressing tumor cell growth. The regulatory mechanisms of this metabolic reprogramming not only help in understanding the metabolic heterogeneity of tumors but also provide a theoretical foundation for developing novel anticancer drugs[39]. Currently, the GLS1 inhibitor CB-839 has entered clinical trial stages[40]. Nrf1 is a critical regulator of intracellular redox homeostasis, and Nrf1α has been identified as an effective tumor suppressor that inhibits cancer progression in xenograft mouse models. Knockout of Nrf1α in human hepatocellular carcinoma cells results in enhanced tumor growth and metastatic potential[41]. Our study revealed that Nrf1α-knockout HepG2 cells exhibited rapid cell death following glutamine deprivation. Additionally, NRF1-knockout HepG2 cells demonstrated increased glycolysis, fatty acid synthesis, glutamine catabolism, and glutamine transport. These findings suggest that Nrf1α gene knockout promotes glutamine dependency。

GLUL (Glutamine Synthetase) is the key enzyme that catalyzes the synthesis of glutamine from glutamate and ammonia[42]. GLUL plays a crucial role in glutamine addiction; its high expression in tumor cells promotes glutamine synthesis to meet the elevated cellular demand for glutamine. Glutamine is converted into glutamate by glutaminase (GLS), which further generates α-ketoglutarate (α-KG) and enters the tricarboxylic acid cycle (TCA cycle), providing carbon and nitrogen sources for cells. High expression of GLUL is closely associated with enhanced glutamine metabolism. For instance, in gliomas, GLUL expression levels positively correlate with tumor malignancy, and knocking out GLUL significantly inhibits tumor cell proliferation and invasion[42-44]. By promoting glutamine metabolism, GLUL supports the energy requirements of tumor cells. The α-KG produced from glutamine metabolism can enter the TCA cycle to replenish metabolic intermediates, thereby maintaining cellular energy metabolism. Targeting GLUL has become a potential strategy for cancer therapy. For example, inhibiting GLUL activity or expression can reduce glutamine synthesis, thereby suppressing tumor cell growth and drug resistance[45].xCT (SLC7A11) is a cystine/glutamate antiporter primarily responsible for transporting extracellular cystine into the cell while exporting glutamate. In cancer cells, high expression of xCT leads to increased intracellular glutamate consumption, which, through a feedback mechanism, prompts cells to absorb more glutamine to replenish glutamate. This mechanism enhances the dependency of cancer cells on glutamine. Significant progress has been made in researching targeted drugs against xCT and glutamine metabolism[46]. By inhibiting xCT’s cystine transport function, the antioxidant defense mechanisms of cancer cells are blocked, inducing lipid peroxidation and ferroptosis[47].Our research found that after Nrf1α knockout, xCT expression increases while GLUL expression decreases, alleviating the massive death of Nrf1α-KO cells caused by glutamine deprivation(Fig.4). Overexpression of the GLUL gene in Nrf1α-KO cells can synthesize large amounts of glutamine from intracellular glutamate, reducing excessive glutamate efflux. The synthesized glutamine helps maintain mitochondrial function, thereby mitigating cell death(Fig.5).In conclusionOur findings provide a valuable theoretical foundation for the development of drugs targeting glutamine addiction.

## Author contributions

R.D. conducted the majority of the experiments, analyzed all data, created all figures, and drafted the manuscript. Q.Z. and S.H.contributed to the experimental design. R.W. participated in the metabolomics experiments and analysis. X.C.,K.L. and D.L. was involved in the intracellular ROS detection. Y.Z. designed and supervised the study, analyzed all data, and assisted in preparing all figures, including those with comics.

## Acknowledgments

We are greatly thankful to Drs. Lu Qiu (Zhengzhou University, Henan, China) and Yonggang Ren (North Sichuan Medical College, Sichuan, China) for their involvement in establishing those knockout cell lines used in this study. We are also thankful to all the present and past members of Prof. Zhang’s laboratory (at Chongqing University, China) for giving critical discussion and invaluable help with this work. This study was funded by the National Natural Science Foundation of China (NSFC, with three project grants 81872336,82473147 and 82073079) awarded to Prof. Yiguo Zhang.

---

**Fig. S1.**
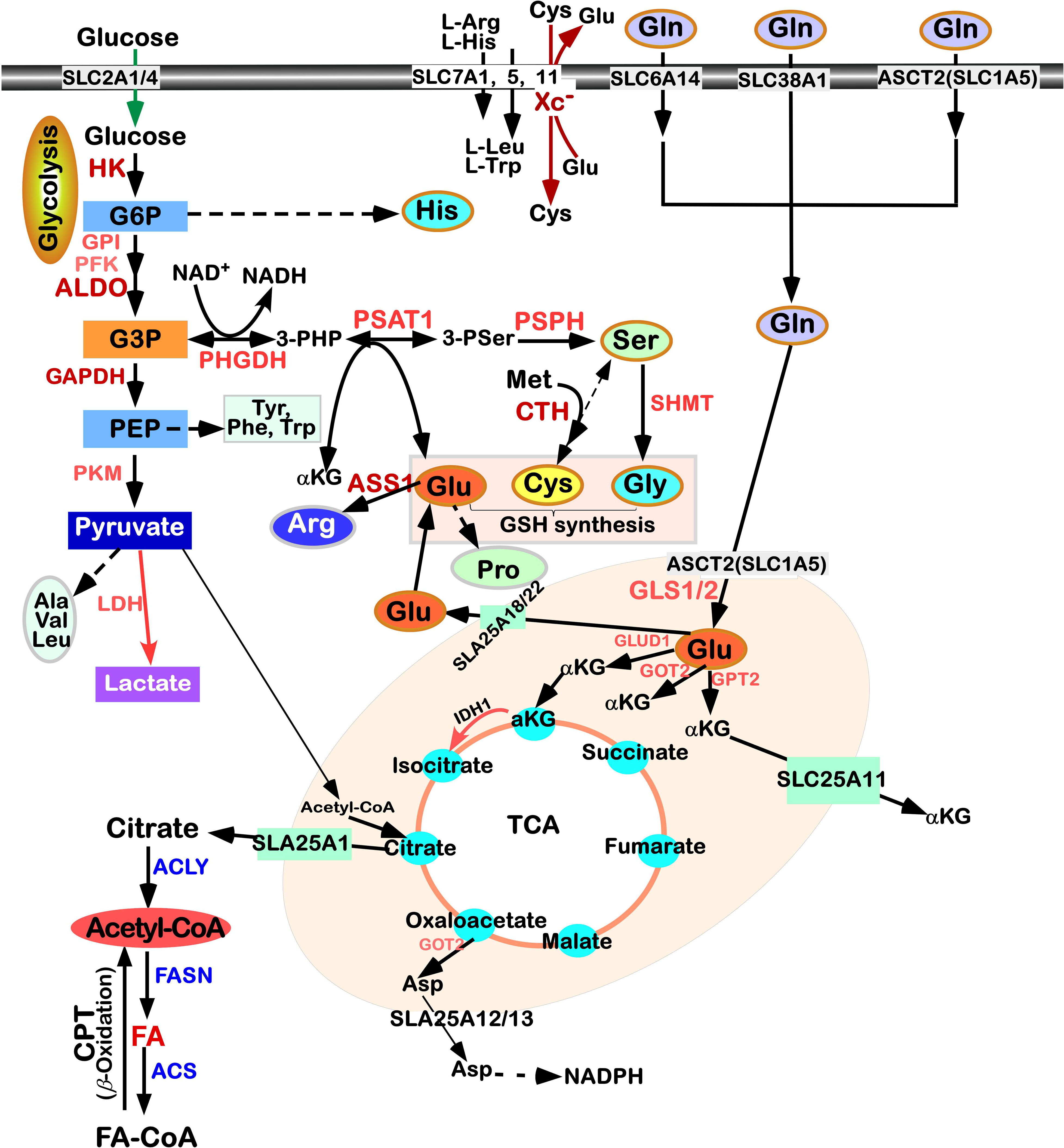
Schematic diagram of cellular glucose metabolism, lipid metabolism, amino acid metabolism, and mitochondrial metabolic processes.

**Fig. S2.**
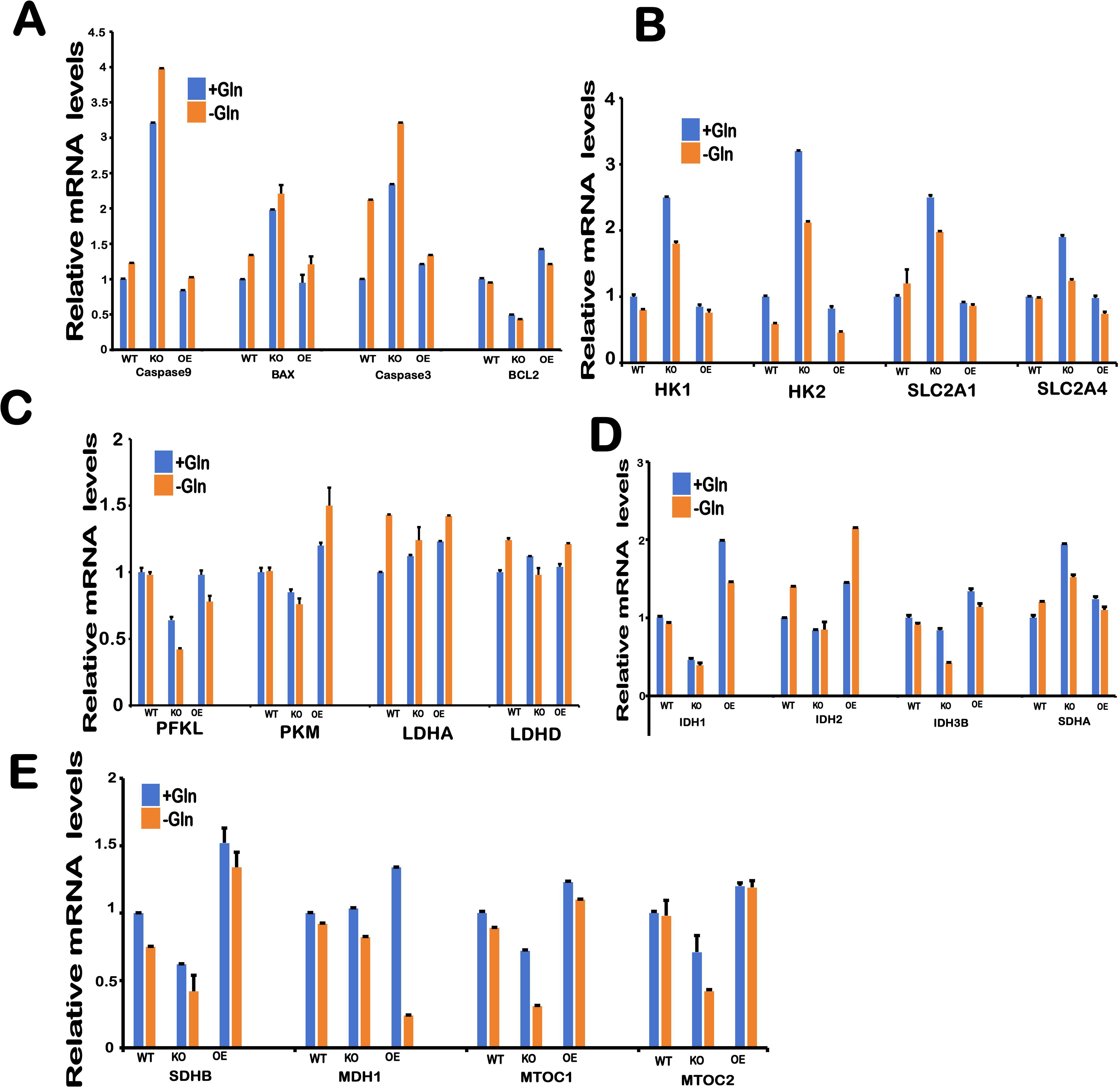
mRNA expression levels of key genes involved in glycolysis and the TCA cycle in WT, Nrf1α-KO, and Nrf1α-OE cells cultured for 24 hours in glutamine-free medium.

**Fig. S3.**
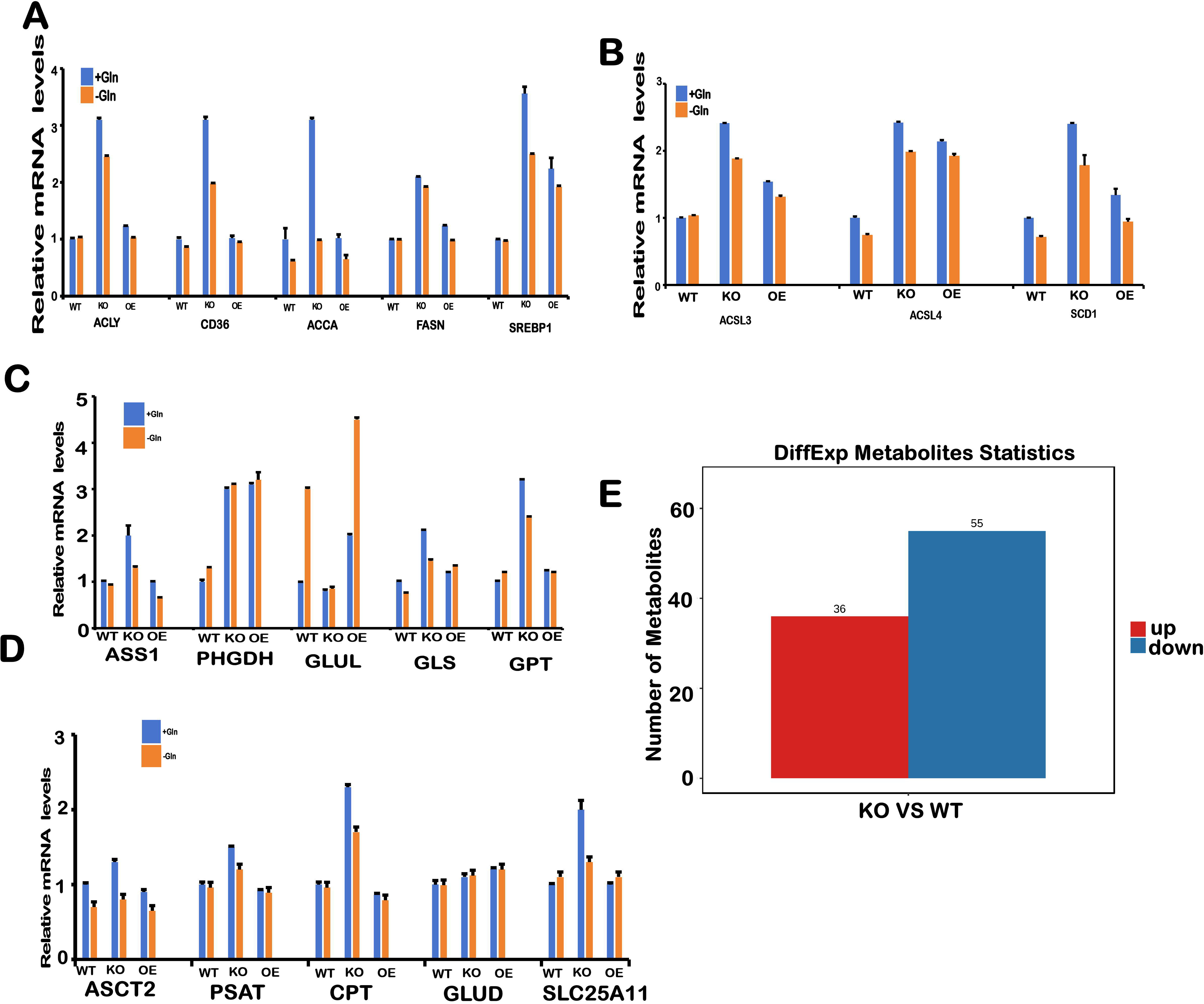
mRNA expression levels of genes related to lipid metabolism and glutamine metabolism in WT, Nrf1α-KO, and Nrf1α-OE cells cultured for 24 hours in glutamine-free medium (panels A-D), and a statistical chart showing differentially expressed metabolites in Nrf1α-KO cells compared to WT cells.

