## Supplemental Table 1 for "Deletion of Nrf1α promotes glutamine addiction of HepG2 cells and thus enhances its apoptosis caused by glutamine deprivation"

**Table 1.**The primer pairs that were used for RT-qPCR analysis

|  | **Forward Primers (5’ to 3’)** | **Reverse Primers (5’ to 3’)** |
| --- | --- | --- |
| Caspase3 | F:ATGGAAGCGAATCAATGGACT | R:ATGGAAGCGAATCAATGGACT |
| Bax | F:CCCGAGAGGTCTTTTTCCGAG | R:CCAGCCCATGATGGTTCTGAT |
| Caspase9 | F:CTCAGACCAGAGATTCGCAAAC | R:GCATTTCCCCTCAAACTCTCAA |
| BCL2 | F:GGTGGGGTCATGTGTGTGG | R:CGGTTCAGGTACTCAGTCATCC |
| Nrf1 | F:AACATTCTGGTCCTTCAGCAATGCT | R:ACCCGTACCCCAATCAAACTCAGCA |
| SLC2A4 | F: TGGGCGGCATGATTTCCTC | R: GCCAGGACATTGTTGACCAG |
| *HK1* | F: CACATGGAGTCCGAGGTTTATG | R: CGTGAATCCCACAGGTAACTTC |
| *HK2* | F: GAGCCACCACTCACCCTACT | R: CCAGGCATTCGGCAATGTG |
| *PFKL* | F: GGCTTCGACACCCGTGTAA | R: CGTCAAACCTCTTGTCATCCA |
| *PKM* | F: ATGTCGAAGCCCCATAGTGAA | R: TGGGTGGTGAATCAATGTCCA |
| *LDHA* | F: ATGGCAACTCTAAAGGATCAGC | R: CCAACCCCAACAACTGTAATCT |
| *LDHD* | F:CCCAGAAGGCAAAGGGAGAG | R: AGGGATTTCTGGCCAGAAGC |
| *IDH1* | F: TGTGGTAGAGATGCAAGGAGA | R: TTGGTGACTTGGTCGTTGGTG |
| *IDH2* | F: TGGCAGTGGTGTCAAGGAGTG | R: GCCCATCGTAGGCTTTCAGTA |
| *IDH3B* | F: GAGCCAAGTCTCAGCGGAT | R: GGGCATCACAAGCACATCAA |
| *SDHA* | F: CAGCATGTGTTACCAAGCTGT | R: GGTGTCGTAGAAATGCCACCT |
| *SDHB* | F: ACCTTCCGAAGATCATGCAGA | R: GTGCAAGCTAGAGTGTTGCCT |
| *MDH1* | F: TTTGGATCACAACCGAGCTAAAG | R: ACATCTGGATACTGAGTCGAGG |
| *MTOC1* | F:ACCCCATTCTATACCAACACCT | ATGGGAGATTATTCCGAAGCCT |
| *MTOC2* | F:GCTGTCCCCACATTAGGCTT | AGATTTCAGAGCATTGACCGTA |
| *ACCα* | F: AACCACATCTTCCTCAACTT | R: ACTTCCATACCGCATTACC |
| *FASN* | F: GTCCACCAGCAACATCAG | R: TTCTCCAGCAAGCCATCT |
| *SREBP1* | F: GGAGCCATGGATTGCACTTT | R: CAGGAAGGCTTCAAGAGAGG |
| *SCD1* | F: CACCACATTCTTCATTGATTGC | R: TCAGCCACTCTTGTAGTTTCCA |
| *Acly* | F: ATCGGTTCAAGTATGCTCGGG | R: GACCAAGTTTTCCACGACGTT |
| *ACSL3* | F: CCACGCAGCGATTCATGAACAT | R: AGCAAACTAATGGTGCTCCCACT |
| *ACSL4* | F: GGTAGAAGGAACTTGGGTTGATA | R: CGTTCAATGTCTTTGAGGTAATG |
| *CPT2* | F: AAGCTGATGAGTAGTGGCAATGA | R: TTGGCGATAATGAGGTTAAAGGA |
| *CD36* | F: CATCGCTGGGGCTGTCATT | R: TGTTGCTGCTGTTCATCATCACT |
| *PHGDH* | F: GAAATCTCTCACGGGGGTTG | R: GTTCACATCCGCCTGCTTG |
| *PSAT* | F: ACCTCAACCCAGATGCCTC | R: TCACGGACAATCACCACGG |
| *ASS1* | F: ATGTCCAGCAAAGGCTCCGT | R: CCAGATGAACTCCTCCACAAACTC |
| *GPT2* | F: GCTCTTTCTCCTGGCTGATG | R: CCTCCTCTGTAACCACACTCG |
| *GLUD* | F: GTGTGTGGAAGAGTTGCCTG | R:CTGCAGGCCTTCGATTGTAC |
| *GLS* | F:TGGAAAAGAGCCGAGTGGAC | R:CCCCCAGCAACTCCAGATTT |
| *SLC25A11* | F:CGCCTTGGCATCTATACCGT | R:CCCTCGTAGCGGACAACTTT |
| *ASCT2* | F:ATCCATGGGCTCCTGGTACT | R:GGGCAGCTCACTCTTCACTT |
| *β-actin* | F: CATGTACGTTGCTATCCAGGC | R: CTCCTTAATGTCACGCACGAT |
