## Supplemental Table 2 for "Deletion of Nrf1α promotes glutamine addiction of HepG2 cells and thus enhances its apoptosis caused by glutamine deprivation"

**Table 2.**The key antibodies used in this work.

| **Antibodies** | **Source** | **Identifier** |
| --- | --- | --- |
| Nrf1 | Zhang’s | N/A |
| Nrf1 | CST | 8052T |
| Caspase3 | CST | 9662S |
| Bax | proteintech | 50599-2-Ig |
| Caspase9 | CST | 9502T |
| BCL2 | proteintech | 12789-1-AP |
| SLC2A1 | ABclonal | A11208 |
| SLC2A4 | ABclonal | A7637 |
| HK1 | proteintech | 19662-1-AP |
| HK2 | Abcam | ab227198 |
| PFKL | Sangon Biotechnology | D222865 |
| PKM | Sangon Biotechnology | D262511 |
| LDHA | ABclonal | A1146 |
| LDHD | ABclonal | A15965 |
| CS | Sangon Biotechnology | D224568 |
| IDH1 | Sangon Biotechnology | D221821 |
| IDH2 | Sangon Biotechnology | D222530 |
| IDH3B | Sangon Biotechnology | D221733 |
| SDHA | Sangon Biotechnology | D223104 |
| SDHB | Sangon Biotechnology | D262175 |
| MDH1 | Sangon Biotechnology | D222668 |
| MT-CO1 | ABclonal | A17889 |
| MT-CO2 | ABclonal | A11913 |
| CD36 | Abcam | Ab133625 |
| ACLY | Sangon Biotechnology | D221957 |
| ACCα | Sangon Biotechnology | D155300 |
| FASN | Sangon Biotechnology | D262701 |
| ACSL3 | Sangon Biotechnology | D261226 |
| ACSL4 | Sangon Biotechnology | D221771 |
| SCD1 | Sangon Biotechnology | D162163 |
| PHGDH | ABclonal | A10461 |
| PSAT1 | ABclonal | A6707 |
| SREBP1 | ABclonal | A26708 |
| SREBP2 | ABclonal | A13049 |
| GLUD | ABclonal | A21792 |
| SLC25A11 | ABclonal | A25022 |
| GLS | ABclonal | A11043 |
| ASCT2 | ABclonal | A23156 |
| GLUL | proteintech | 66323-1-Ig |
| CPT | proteintech | 26555-1-AP |
| GPT2 | ABclonal | A11819 |
| ASS1 | Sangon Biotechnology | D224199 |
| β-actin | ZSGB-BIO | TA-09 |
